# Injury and a program of fetal wound healing in the fetal and neonatal extrahepatic bile duct

**DOI:** 10.1101/2022.12.15.520656

**Authors:** Iris E.M. de Jong, Mallory L. Hunt, Dongning Chen, Yu Du, Jessica Llewellyn, Kapish Gupta, Dorothea Erxleben, Felipe Rivas, Adam R. Hall, Emma E. Furth, Ali Naji, Chengyang Liu, Abhishek Dhand, Jason Burdick, Marcus G. Davey, Alan W. Flake, Robert J. Porte, Pierre A. Russo, J. William Gaynor, Rebecca G. Wells

## Abstract

**Introduction:** Biliary atresia (BA) is an obstructive cholangiopathy that initially affects the extrahepatic bile ducts (EHBDs) of neonates. The etiology is uncertain, but evidence points to a prenatal cause; however, the response of the fetal EHBD to injury remains unknown. The objective of this study was to define the fetal response to EHBD injury and to determine whether it follows a fetal wound healing paradigm.

**Methods:** Mouse, rat, sheep, and human EHBD samples were studied at different developmental time points. Models included a fetal sheep model of prenatal hypoxia, human BA EHBD remnants and liver samples taken at the time of the Kasai procedure, EHBDs isolated from neonatal rats and mice, and spheroids and other models generated from primary neonatal mouse cholangiocytes.

**Results:** A wide layer of high molecular weight HA encircling the lumen was characteristic of the normal perinatal but not adult EHBD. This layer, which was surrounded by collagen, expanded in injured ducts in parallel with extensive peribiliary gland (PBG) hyperplasia, increased mucus production and elevated serum bilirubin levels. BA EHBD remnants similarly showed increased HA centered around ductular structures compared with age-appropriate controls. High molecular weight HA typical of the fetal/neonatal ducts caused increased cholangiocyte spheroid growth, whereas low molecular weight HA induced abnormal epithelial morphology; low molecular weight HA caused matrix swelling in a bile duct-on-a-chip device.

**Conclusion:** The fetal/neonatal EHBD, including in human EHBD remnants from Kasai surgeries, demonstrated an injury response with high levels of HA typical of the regenerative, scarless program termed fetal wound healing. Although generally beneficial, the expanded peri-luminal HA layer may swell and lead to elevated bilirubin levels and obstruction of the EHBD.

## INTRODUCTION

Biliary atresia (BA) is a devastating disease of newborns primarily affecting the extrahepatic bile ducts (EHBDs). Patients with BA are born seemingly healthy but develop a (sub)total obstruction of the EHBD days to weeks after birth followed by the rapid development of hepatobiliary fibrosis and cirrhosis. Although the etiology of BA remains under investigation, compelling evidence points to a prenatal environmental insult [1,2].

Mechanisms of fetal EHBD repair are not known. Repair of the injured adult EHBD requires peribiliary glands (PBGs), which are mucus-producing glands in the submucosa that harbor a cholangiocyte stem/progenitor cell population. With severe damage to the duct, these cells proliferate, migrate towards denuded luminal surfaces, and differentiate to mature cholangiocytes [3,4]. Additionally, there may be associated extracellular matrix (ECM) deposition and scarring or fibrosis, as per a typical adult wound healing program.

Healing during much of the fetal period, however, is scarless, as demonstrated in multiple organs including heart, skin, lung, and tendon [5–8]. Although not previously studied in the EHBD, fetal wound healing in general is dominated by deposition of the glycosaminoglycan hyaluronic acid (HA) rather than the type I collagen typical of adult wound healing [9]. HA is a highly-charged polymer consisting of numerous repeats of D-glucuronic acid and N-acetylglucosamine and is a component of the normal ECM of the EHBDs at all stages of development, at least in mice [10]. HA is synthesized in high molecular weight form (HMW, >1MDa) by hyaluronic acid synthases (HAS) and degraded from a high to a low molecular weight (LMW) form by hyaluronidases (HYAL); a dynamic process of production and degradation regulates the average molecular mass of HA in tissues [11,12]. HMW HA is critical in embryogenesis, facilitating proliferation, cell migration, growth, and development. HA accumulation during fetal wound healing occurs typically in HMW form and is implicated in scarless, regenerative wound healing. In contrast, LMW HA (<250kDa) is associated with pro-inflammatory properties and increased fragmentation and is typical of pathological conditions including adult wound healing.

We hypothesized that the injured fetal EHBD follows a program of fetal wound healing. The goal of this study was to define the fetal response to EHBD injury and in particular to determine whether fetal EHBD repair follows the fetal wound healing paradigm and whether this has implications for the development of BA.

## METHODS AND MATERIALS

### Animals

Handling and care of rodents was carried out according to protocols that were approved by the University of Pennsylvania Institutional Care and Use Committee, as per the National Institutes of Health Guide for the Care and Use of Laboratory Animals (protocol 804862). Animals were selected for use in experiments by age, regardless of sex. EHBDs were dissected from the distal end just before entering the pancreas to the proximal end just below the cystic duct. EHBD and bile samples from sheep were collected as excess tissue as part of previously described experiments whereby fetuses were maintained in utero (“normal”, 20-25 ml/kg/min oxygen) or in an external biobag and supplied with 14-16 ml/kg/min oxygen (“hypoxia”) [13,14]. The handling and care of sheep complied with the ethical standards of the National Institutes of Health Guide for the Care and Use of Animals and protocols were approved by the Institutional Animal Care and Use Committee (IACUC) of The Children’s Hospital of Philadelphia (protocol 1212).

### Human samples

Anonymized human adult EHBD sections were collected as part of the Human Pancreas Procurement and Analysis Program (HPPAP), which was granted exemption from the University of Pennsylvania Institutional Review Board (protocol 826489). Subjects (7 adults, 23-67 years old) were otherwise healthy individuals who died unexpectedly, with consent obtained from next of kin. De-identified, formalin-fixed paraffin embedded (FFPE) human EHBD tissue sections from 4 fetuses (22, 34, 36, and 39 weeks of gestation), 3 infants (2, 5, and 6 months old), and 16 BA remnants derived from the porta hepatis were obtained from the Anatomic Pathology archives of the Children’s Hospital of Philadelphia; all were excess tissue from samples obtained for clinical indications and were IRB exempt. Of the BA remnant sections, 9 included epithelial structures. The study also included FFPE samples that corresponded to the large intrahepatic bile ducts from 3 adults with primary sclerosing cholangitis (PSC), obtained from the pathology archives of the University of Pennsylvania (IRB protocol 831726).

Further details regarding the materials and methods are reported in supplementary information.

## RESULTS

### Fetal and neonatal EHBDs have higher HA than adult ducts, localized in a periluminal ring

We first determined the differences in HA and collagen composition between the fetal and adult EHBD. As has been reported for other organs, staining using HA binding protein (HABP) showed that HA in the EHBD is high in the fetus and neonate compared to the adult [15] across a range of species (humans, sheep, rats, and mice) (**Figures 1A and S1**). When we quantified EHBD HA content biochemically, we found a decrease in percent HA content from neonates to adults (**Figure 1B**). This was accompanied by a gradual increase in content of type I collagen (**Figure S2**). Surprisingly, HA in fetal and neonatal EHBDs was almost entirely located in a dense layer directly around the lumen, while collagen was peripheral; as development progressed, HA around the lumen gradually decreased and was replaced by collagen, which, as we previously reported and also observed here, was deposited from the outside in (**Figures 1A,D and S2**) [10]. Accordingly, the ratio of collagen to HA increased over time (**Figure 1C**). Second harmonic generation (SHG) imaging and Sirius Red staining for collagen showed a clear distinction between the HA layer and the more peripheral collagen layer in the fetus, especially in the larger species (human/sheep) (**Figure 1D,E**). Note that the human fetal EHBD sample was obtained at a gestational age of 36 weeks, whereas the sheep fetus EHBD sample corresponded to a gestational age of 121 days (mean gestation in sheep is ~147 days) (**Figures 1D,E**). Much of the embryological development of the EHBD is completed after birth in rodents, comparable with the last trimester in human [16,17]. This is in line with our HA staining showing a wide layer of HA in the EHBD wall on postnatal day 1 in rodents resembling the HA staining of the fetus sheep and human EHBD (**Figure 1A**).

**Figure 1.**
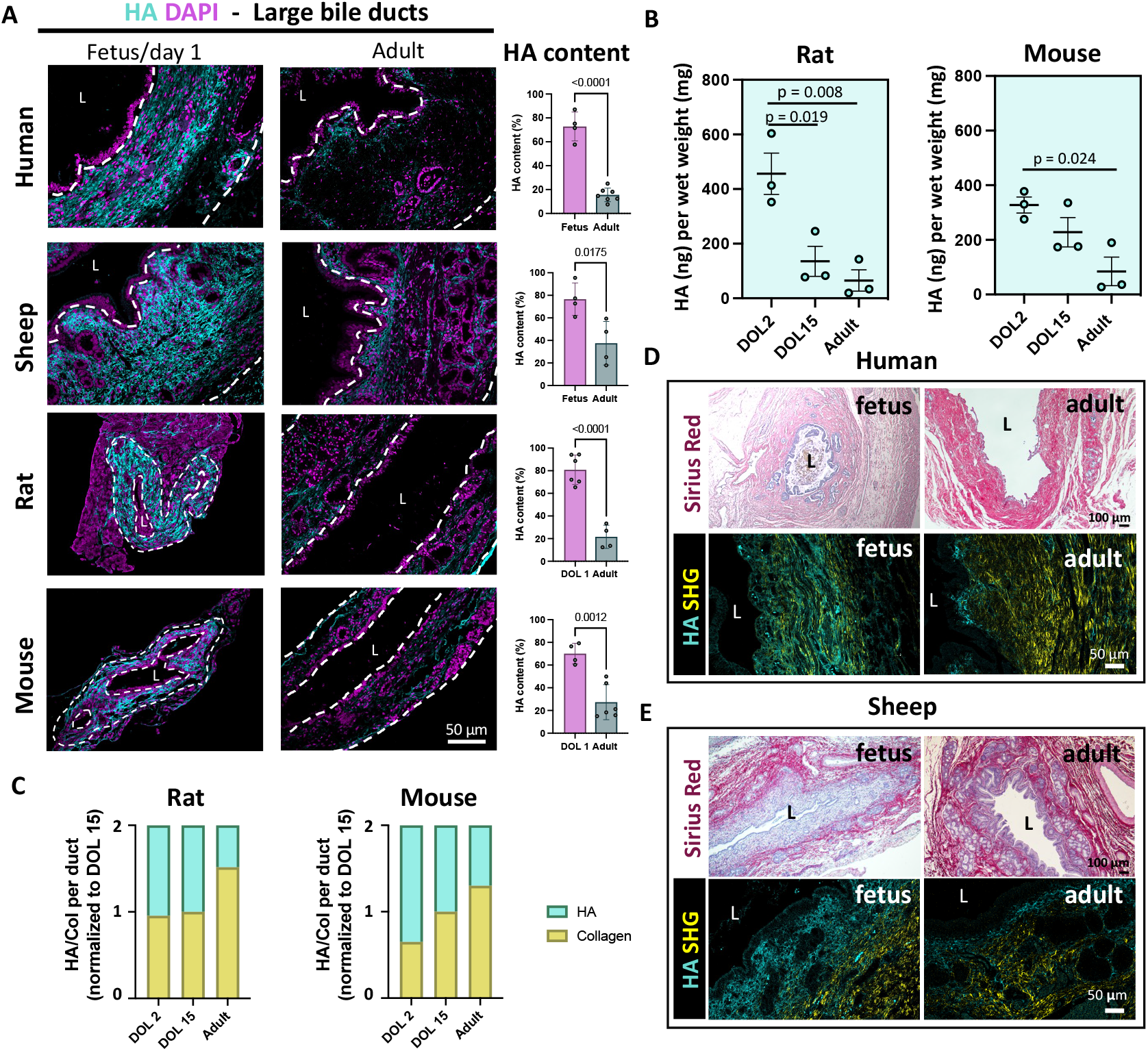
Normal mammalian fetal and neonatal EHBDs have high HA and low collagen compared to the adult. **(A)** HA content of adult or fetal/neonatal human (GA: 22-39 weeks), sheep (GA: 121 and 128 days), and rat and mouse EHBDs (DOL 1). HA content was defined as % coverage of wall by HA binding protein signal (n ≥ 3 individuals). **(B)** Concentration of HA (ng) per mg wet weight EHBD. n = 3 homogenates, each containing 2-15 EHBDs. **(C)** Ratio of HA and collagen in the EHBD, normalized to DOL15. **(D)** Human fetus (GA: 36 weeks) and adult EHBD. SHG imaging includes both forward and backward scatter **(E)** Sheep fetus (GA: 121 days) and adult EHBD. All graphs show mean±SD. Significance determined by Student’s t test (A) or One-Way ANOVA with Tukey’s post-hoc test (B).

Taken together, these data show that the ECM content and distribution in fetal/neonatal EHBDs is fundamentally different from the adult EHBD, providing a substantially different environment – HA predominant in the fetus, collagen in the adult – for bile duct regeneration.

### Prenatal EHBD damage increases HA

HA is a major mediator of scarless wound healing in the fetus [9]. We therefore asked whether there are changes in the HA layer after EHBD damage. To study fetal EHBD wound healing, we used a sheep model of extra-uterine gestation (**Figure S3A**). After 14-21 days under hypoxic conditions, there were significant differences between the EHBDs of sheep maintained ex utero compared to sheep of the same gestational age that remained in utero. This included patchy detachment of the surface epithelium with mural bleeding and a statistically significant expansion in the width of the HA layer directly around the lumen (**Figures 2A, top row and S3B**). We observed a similar significant increase in the thickness of the peri-luminal HA layer in biliary remnants removed at the time of Kasai hepatoportoenterostomy from patients with BA, as compared to controls comparable in age (2-6 months of age) (**Figure 2A, middle row)**. In contrast, large bile ducts of adults with primary sclerosing cholangitis (PSC) did not show a significant expansion of the HA layer compared to adult controls (**Figure 2A, lower row**). Viewed in another way, the distance between the lumen and the collagen layer appeared greater in the injured fetal sheep EHBDs and human BA remnants compared to age-appropriate controls (**Figures 2B and S4A,B**). Although the BA remnant samples typically lacked a lumen, any epithelial structures visualized were surrounded by HA, with collagen outside of the HA ring (**Figures 2B, 3A, and S4A,C**). The main HA-producing enzymes are hyaluronic acid synthases (HAS) 1, 2, and 3 [9]. Interestingly, we observed an accumulation of HAS1-3-expressing mesenchymal cells near the lumen in fetal sheep EHBDs, mostly after hypoxia (**Figures S5A,B**).

**Figure 2.**
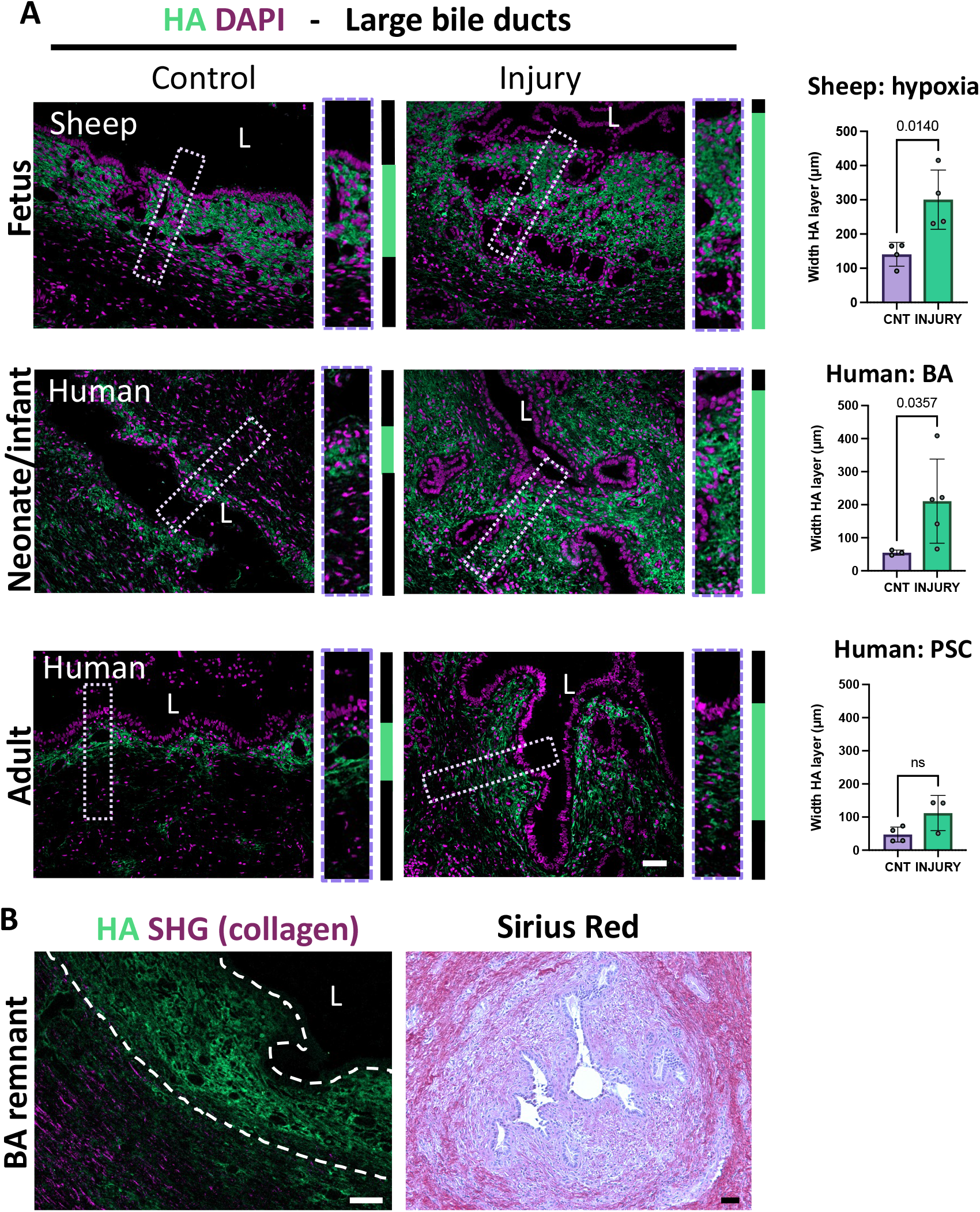
Prenatal EHBD injury causes an increase in HA. **(A)** HA layer of damaged/diseased and normal fetal sheep (GA: 128 days) and infant and adult human EHBDs. Significance determined by Student’s t-test (fetal sheep) and Mann-Whitney test (infant and adult human). **(B)** Representative images of n = 9 EHBD BA remnants. Overlay of immunohistochemistry for HA (green) and visualization of collagen by SHG imaging (forward and backward scatter are both magenta) and Sirius Red staining. Graphs show mean±SD of n ≥ 3 individuals for each condition. All scale bars = 50 μm.

### BA remnant collagen fibers differ from controls

Given that that HA can disrupt collagen organization [18], we asked whether there were organizational differences between the collagen in EHBD remnants from BA patients and controls. The structure of the collagen around the HA layers in BA remnants was determined using the ImageJ plugin TWOMBLI to identify matrix patterns. It showed that collagen fibers in Kasai remnants from BA patients were shorter and had more variable curvature, but did not differ in density compared to controls (**Figures 3A-E**). This is consistent with the collagen organization that is observed in (adult) pathological processes [19].

**Figure 3.**
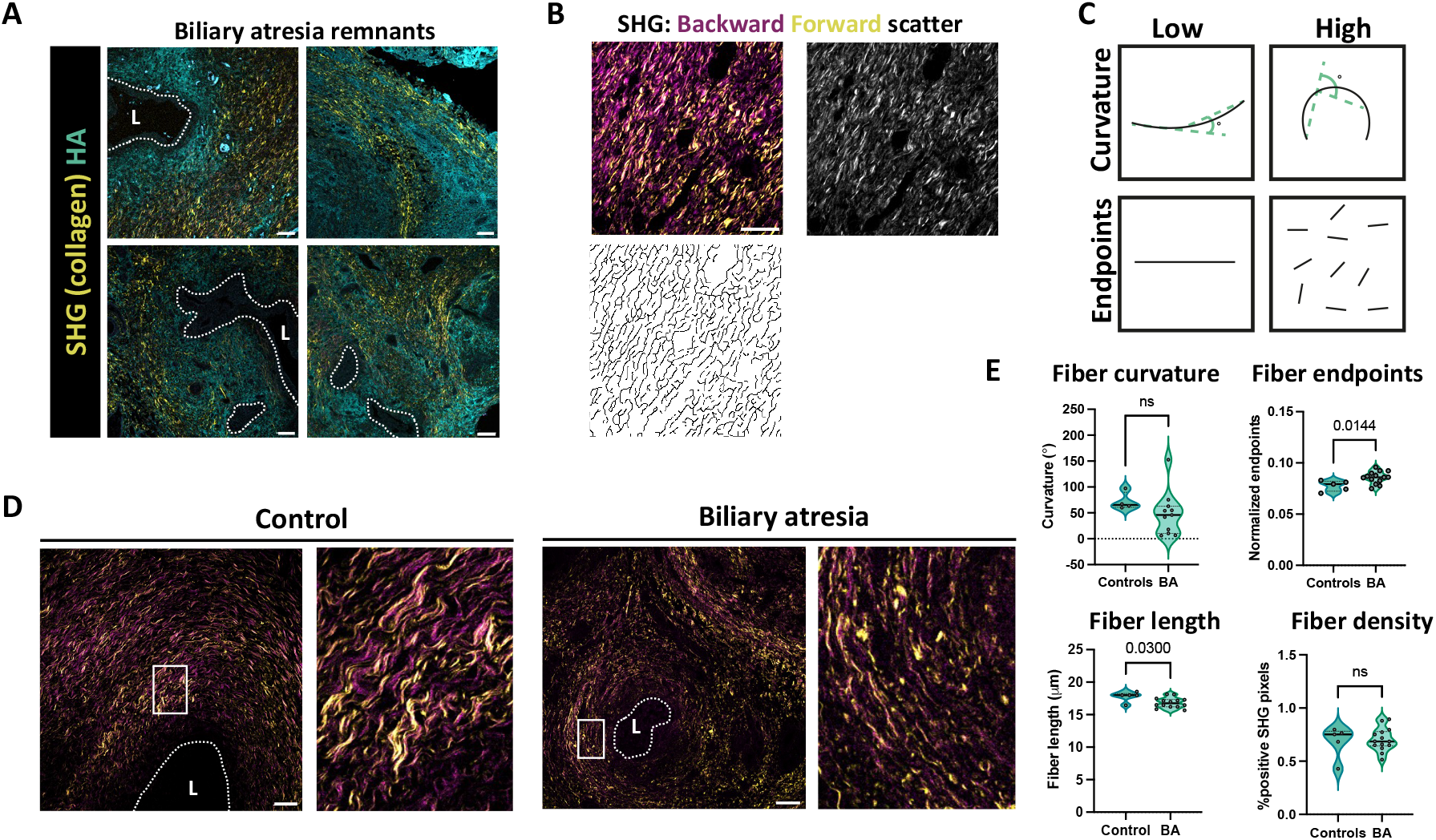
Collagen in BA is organized around HA areas and the fibers are shorter. **(A)** Representative images of n = 14 BA remnants, with collagen visualized by SHG imaging and HA by HA binding protein. **(B)** Mask rendering from SHG signals (both forward and backward scatter) on a control sample (age: 2 months) **(C)** Schematic of curvature and endpoint calculations. **(D)** Representative SHG imaging of EHBDs from human controls (n = 5; age: 2-6 months) and BA remnants (n = 14) taken at the Kasai procedure. **(E)** Fiber curvature, endpoints, length, and density. Significance determined by Student’s t test. All scale bars = 50 μm.

### Fetal EHBD injury leads to a robust progenitor cell response

We hypothesized that the differences between the ECM of the normal adult and the fetal EHBD reflect different programs of wound healing. Epithelial regeneration after injury of the adult EHBD has been described in the context of adult cholangiopathies [20]. The injured EHBDs of fetal sheep showed marked regeneration with the presence of a double layered surface epithelium, PBG hyperplasia (i.e., increased PBG area), and increased proliferation of epithelial and mesenchymal cells (**Figures 4A,B**). PBGs were distributed regularly throughout the submucosa in both the control and the hypoxia group in fetal EHBDs (**Figure 4A**), in contrast to adult EHBDs in which the PBGs generally appear in compact clusters [3]. In fetal sheep incubated in the external womb, the area occupied by PBGs expanded in proportion to the duration of time in hypoxic conditions, and increased in proportion to the increase in width of the EHBD HA layer (**Figures 4C-E**).

**Figure 4.**
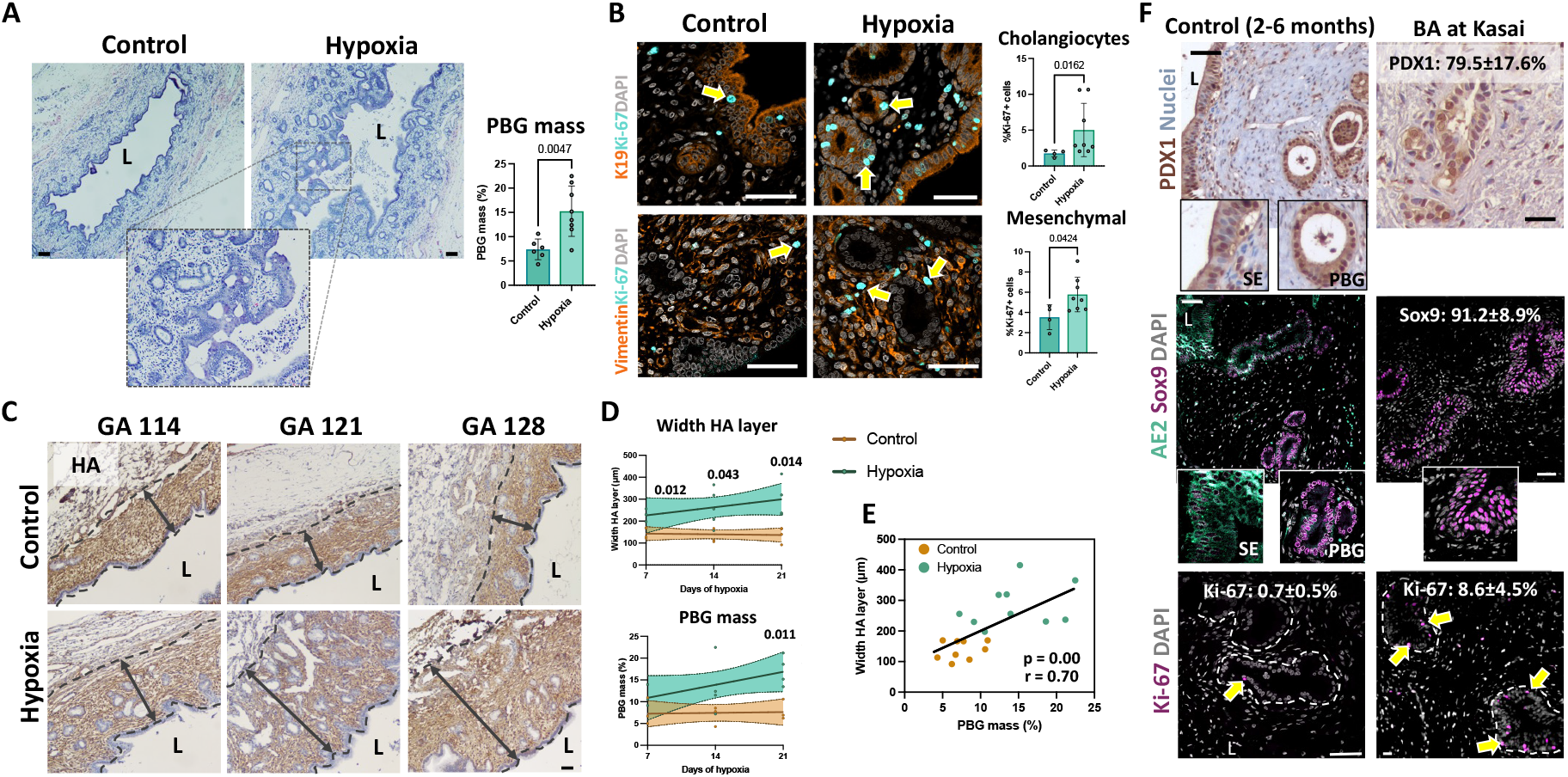
Prenatal hypoxia causes a marked epithelial regenerative response in EHBDs of fetal sheep. **(A)** H&E staining of EHBDs from control (n = 6) and hypoxia (n = 8) groups (GA: 121 and 128 days) and graphs representing PBG area. **(B)** Immunofluorescence for Ki-67 (proliferation), K19 (cholangiocytes), and vimentin (mesenchymal cells) on sheep fetus EHBDs (GA 121 and 128). n = 4 (controls) and n = 8 (hypoxia). **(C)** Representative HA stains (brown) in control and hypoxia groups (n ≥ 3 for each timepoint). **(D)** HA layer width (from C) and PBG area (from A). Graphs show mean±SE of the best curve to fit the data. **(E)** Correlation between the HA layer width and PBG area (as plotted in D). **(F)** Human control EHBDs (n = 5; age: 2-6 months) and BA remnants (n = 14) stained for PDX1, AE2, Sox9, and Ki-67. %positive cells is provided. Significance determined by Student’s t-test (A-D), Mann-Whitney (B:cholangiocytes) or Spearman correlation (E). Graph shows mean±SD (all panels). All scale bars = 50 μm.

High percentages of Sox9+ and PDX1+ cholangiocytes were observed in the BA remnants, resembling the PBGs of the control neonatal/infant (age: 2-6 months) EHBDs (~100% of the PBG cells were PDX1+ and Sox9+) (**Figure 4F**). Anion Exchanger 2 (AE2) marks mature and functional cholangiocytes and was generally located in the surface epithelium of control EHBDs, but no cholangiocytes in the BA remnants expressed AE2 (**Figure 4F, middle panel**). The surviving cholangiocytes in BA remnants showed high proliferation rates compared to controls (p < 0.01) (**Figure 4F, lower panel**). In fetal sheep EHBD samples, there was also marked expression of Sox9 although in PBGs as well as in the surface epithelium, suggesting that in fetal life the surface epithelium is still immature (**Figure S6**).

Thus, the fetal biliary regenerative response is distinct from that of the adult [3] and is characterized by marked PBG hyperplasia, marked proliferation, and increased HA (rather than collagen) deposition immediately around the lumen.

### HA deposition by the fetal EHBD after injury predisposes the duct to obstruction

The width of the peri-luminal HA layer as well as the water-retaining (and therefore swelling) qualities of HA led us to hypothesize that the HA deposition in the fetal ducts predisposed to duct obstruction [21]. We therefore examined the fetal sheep in the hypoxia group for markers of biliary obstruction. First, we found that all liver samples taken from the animals in the hypoxia group (n = 11) showed bile plugs in the intrahepatic bile ducts while none of those from the control group had plugs. Additionally, serum bilirubin was significantly increased in the hypoxia group (**Figure 5A**). Second, we measured the density and width of the outer collagen layer in control vs. injured EHBDs and showed that it was narrower in injured ducts compared to controls, and that there was a trend towards increased collagen density, consistent with compression of the collagen bundles by an expanding HA layer (**Figure 5B**). We next modeled the impact of HA in a microfluidic device whereby collagen or collagen plus HMW or LMW HA was gelled around a needle, forming a channel (**Figure 5C**). At baseline after needle removal, the diameters of the channels ranged between 97 – 144 μm and were comparable between the three conditions. We then added PBS to the system and determined changes in lumen diameter. The HA-filled ECM swelled, leading to a decrease in lumen diameter, particularly in the LMW HA-containing device (**Figures 5C, D**).

**Figure 5.**
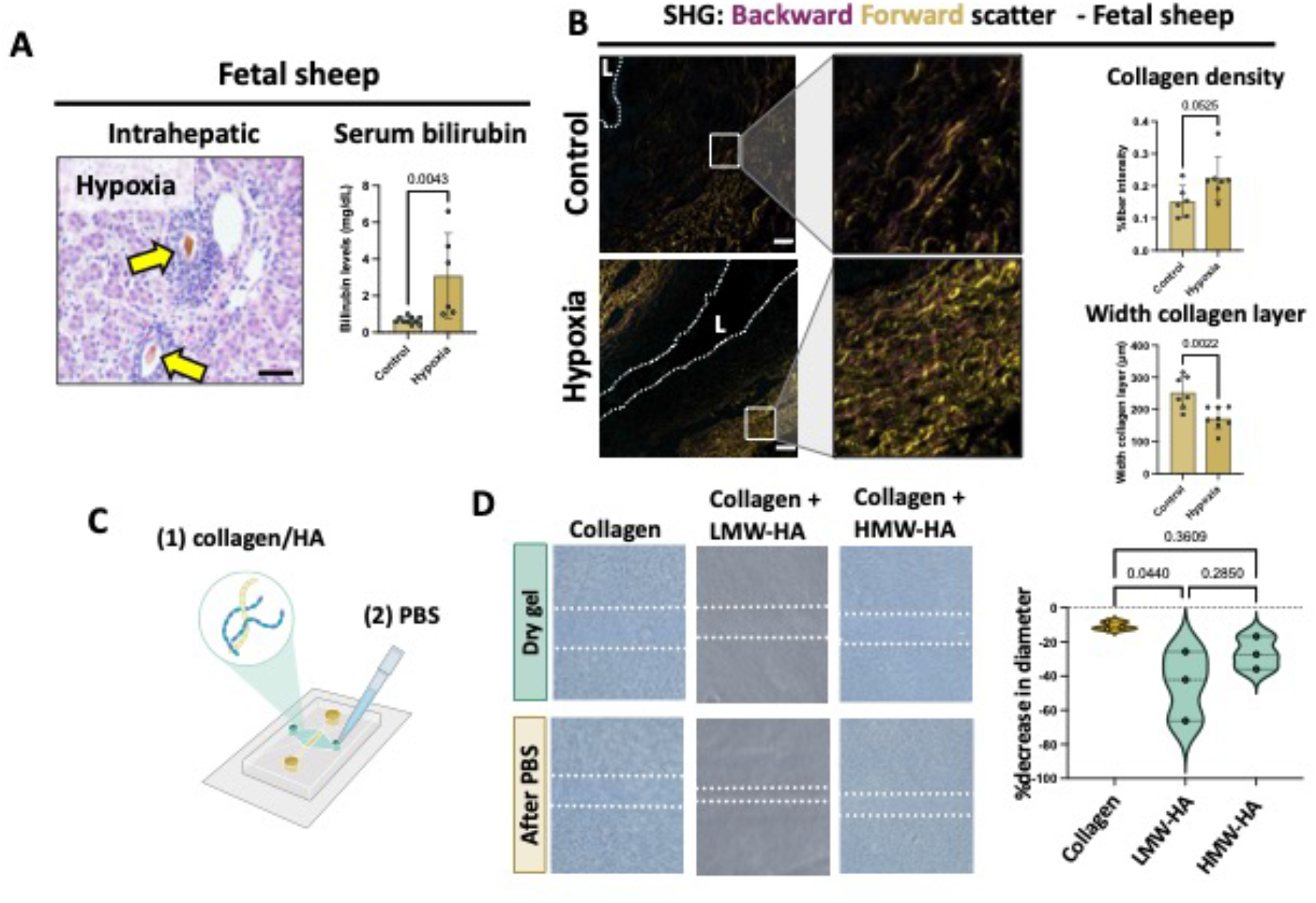
HA deposition predisposes to obstruction. **(A)** H&E staining of a representative fetal sheep liver in the hypoxia group showing bile plugs. Serum bilirubin levels of fetuses in the control (n = 10) and hypoxia (n = 6) groups (GA: 121 and 128 days). **(B)** SHG imaging of EHBDs from control (n = 7) and hypoxic fetal sheep (n = 8) to determine collagen density and width of collagen layers. **(C)** Setup of the microfluidic device. **(D)**. (left) Devices with the three types of matrices (n = 3, each), before and after swelling. (right) Violin plot of % change in lumen diameter for each group. Significance determined by Student’s t-test or One-Way ANOVA with Tukey’s post-hoc test (D). Graphs show mean±SD. All scale bars = 50 μm.

Thus, there is compelling evidence for obstruction in the ducts of injured fetal sheep, even in the absence of liver or duct fibrosis; and in vitro modeling as well as the matrix structure of the EHBDs are consistent with HA-associated swelling as a cause.

### EHBD regeneration in fetal sheep promotes mucus production

We also considered the properties of bile after fetal EHBD injury. In addition to there being increased numbers of PBGs in EHBDs from hypoxic fetal sheep (**Figures 4A,D**), the PAS-positive area of the PBGs was consistently higher in the hypoxia group compared to controls, indicating increased mucin production (**Figure 6A**). In the hypoxia group, mucin-positive cells were evident in both PBGs and the surface epithelium, which contained goblet cells (**Figures 6B and S7A**), in line with reports of goblets cells in human fetal EHBDs in later gestation [22]. Mucus produced by PBGs, like all mucus, has a high viscosity and contributes to the viscosity of bile [23]. Based on the number of mucin-containing cells in EHBDs in the hypoxia group, we hypothesized that the viscosity of bile from these animals would be increased. However, there was no bile apparent in the gallbladders of the hypoxic animals. We therefore compared viscosity of bile samples from the control fetal sheep with bile samples from two adult sheep. Although bile viscosity was higher in a subset of the fetal samples, we were unable to collect enough to determine significance (**Figure S7B**). We hypothesized, however, that if mucus-containing bile accumulated in the EHBD and led to increased obstruction, we would be able to detect mucus in the intrahepatic bile plugs.

**Figure 6.**
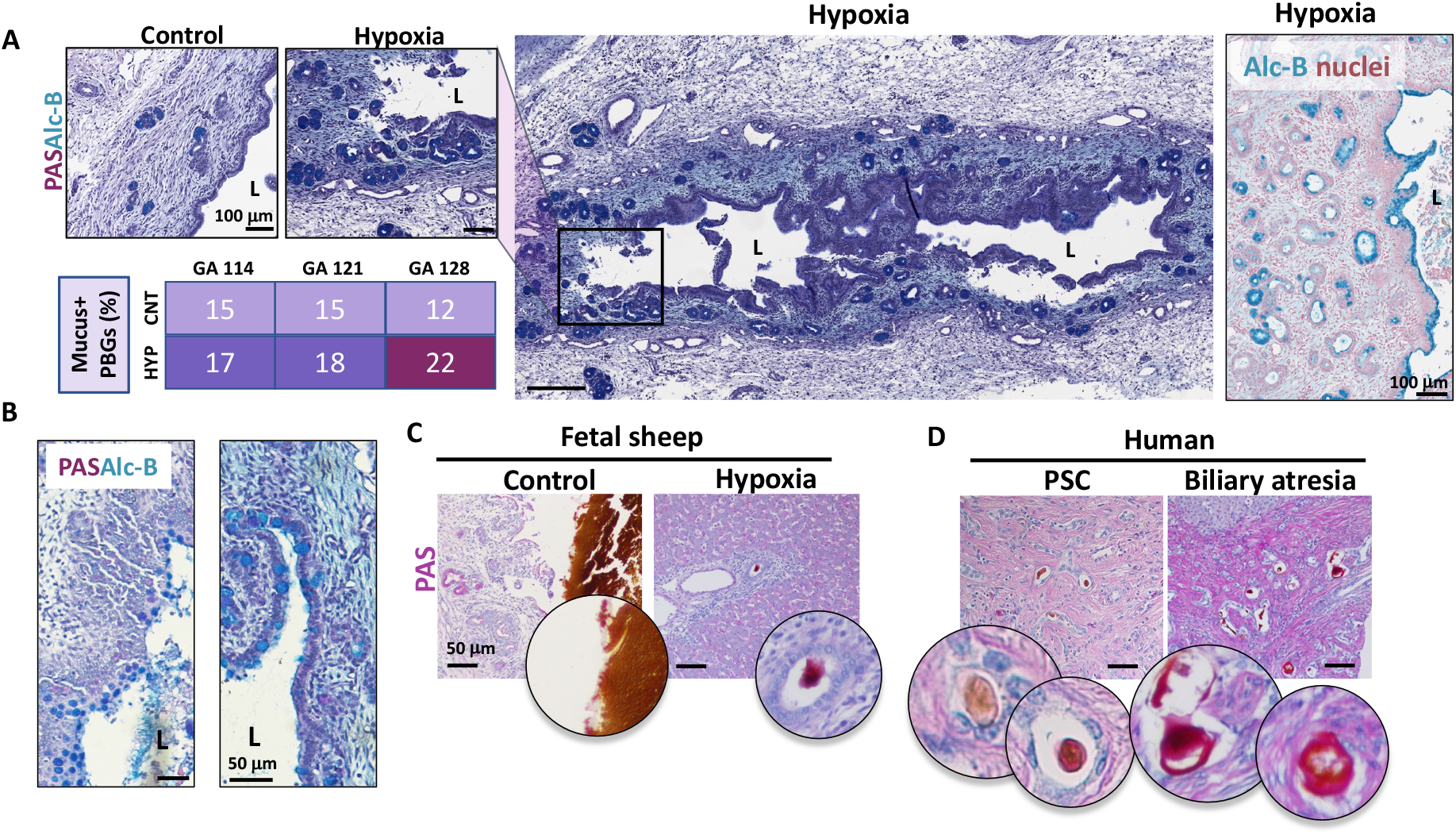
The fetal EHBD regenerative response involves an increase in mucus production. **(A)** PAS and alcian blue-stained EHBDs from fetal sheep in control and hypoxia groups. Table indicates % PBGs showing mucus production for the indicated gestational age (n ≥ 3 for each timepoint). Colors, from light to dark purple, correspond to increasing percentages (<15%, <20%, and >20%). **(B)** Combined PAS and alcian blue staining of goblet cells in the surface epithelium of the hypoxia group. **(C)** PAS staining of bile in the gallbladder (controls did not show intrahepatic bile plugs) and bile plug in the intrahepatic bile ducts **(D)** PAS staining of intrahepatic bile plugs of patients with PSC or BA (n = 3, each).

Bile plugs were only seen in the hypoxia group; they stained positive for PAS, indicating the presence of mucus, in contrast to bile in the gallbladder of controls (**Figure 6C**). Pretreating livers with bilirubin oxidase to minimize potential interference from bilirubin pigments in bile confirmed the positive staining (**Figure S8)**. Similar treatment of livers from patients with BA also showed mucus staining of bile plugs. Normal human livers do not contain bile plugs so cannot be used for comparison, but liver sections from PSC patients showed less prominent mucus in bile plugs than did BA livers (**Figure 6D**).

Collectively, these results suggest that PBG hyperplasia and mucus production, part of the regenerative response of the fetal EHBD, may result in increased bile viscosity, potentially contributing to EHBD obstruction. The damage-repair response of fetal EHBDs includes expansion of the HA layer, which swells, compressing both the collagen layer and the lumen. The decreasing diameter of the lumen may impede bile flow, especially if the bile is already viscous.

### Neonatal EHBDs contain higher molecular weight HA than adults, which is degraded after injury

In order to better understand the dynamics of HA during development and after injury, we studied the expression of the enzymes that synthesize and degrade HA. Neonatal mesenchymal cells showed increased expression of HAS1-3 compared to adults (p = 0.01; p = 0.01; p < 0.01, respectively), indicating that they actively produce HA (**Figure 7A**). These results suggest that production of HA in the EHBD submucosa of neonates is higher than in adults, consistent with our data showing that neonatal ducts have increased HA compared to adults, including on a per weight basis, across species (**Figure 1**). Staining for the two major HA degradation enzymes, HYAL1 and HYAL2, similarly demonstrated higher expression in neonates compared to adults (p < 0.01; p < 0.01) (**Figure 7A**). We therefore used solid-state nanopore technology [24] to determine the size of HA polymers in neonates and adults, and found that HA molecular weight was significantly higher in neonates compared to adults, consistent with an ongoing process of degradation after synthesis (**Figure 7B, Table S2**).

**Figure 7.**
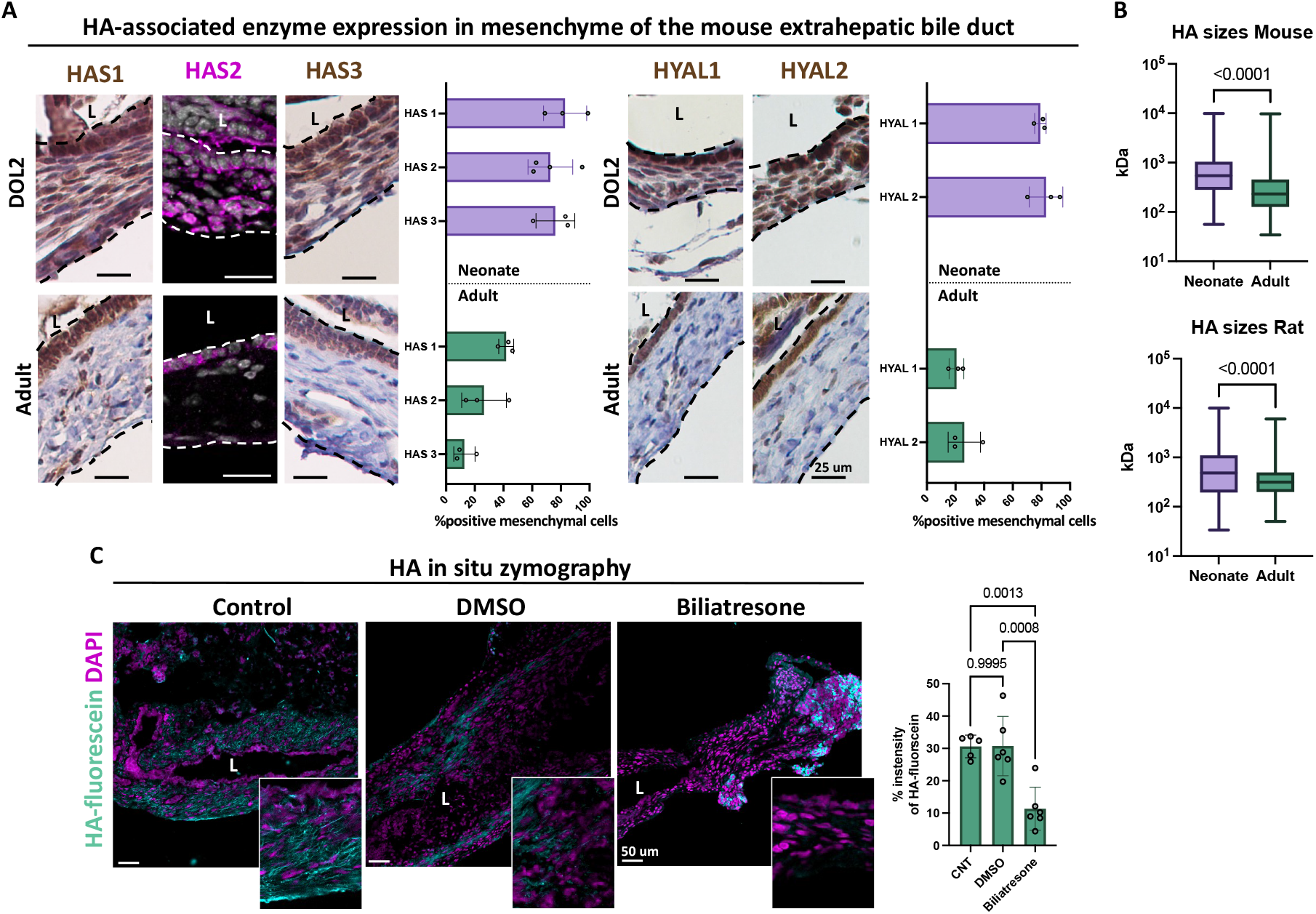
HA size distribution and production/degradation enzymes differ between neonates, adults, and after injury. **(A)** Mouse neonatal EHBDs at day 2 after birth and adult EHBDs stained for HAS1-3 and HYAL1-2. Percentage positive mesenchymal cells for n=3 for each stain shown in graph. **(B)** HA size distribution in mouse and rat neonatal and adult EHBDs, as measured by solid-state nanopore technology. **(C)** HA in situ zymography using fluorescein-labelled HA, overlaid on 4/5 day-old rat neonatal EHBDs after immediate harvest (control; n = 5) or after an additional 5-day treatment with DMSO (n = 6) or biliatresone (n = 6). Significance determined by Student’s t-test or One-Way ANOVA with Tukey’s post-hoc test (C). Graphs show mean ± SD (all panels).

Next, we asked whether injury to neonatal EHBDs would increase HA degradation. Treatment of neonatal rat EHBD explants with biliatresone, a toxin that causes a BA-like phenotype in animal models and in explants [25], resulted in increased breakdown of fluorescein-labelled HA (**Figures 7C** and **S9**). There was some damage in the DMSO-treated EHBDs, likely the result of the explant culture system. Taken together with Figures 1 and 2, these results indicate that production, degradation, and average molecular weight of HA in the submucosa of neonatal bile ducts is higher than in adult ducts; and that additionally, injury leads to both increased production and degradation of HA.

### The size of HA influences EHBD cell behavior in 2D and 3D cultures

Our EHBD data are consistent with extensive literature reporting that adult wound healing is associated with an increased degradation of HA, yielding a high concentration of LMW HA, whereas fetal wound healing is characterized by an accumulation of HMW HA [9,12,26]. To determine the cellular response of the EHBD to LMW and HMW HA, we co-cultured primary neonatal mouse cholangiocytes and EHBD mesenchymal cells in spheroids. Cells were embedded in droplets containing a mixture of Matrigel/collagen or Matrigel/collagen/HA. Cells cultured in mixtures with HMW HA demonstrated increased spheroid growth, diameter, and proliferation rates compared to cells in mixtures with LMW HA (**Figures 8A,B**). Epithelial architecture differed between the groups – in LMW HA, 19% of the spheroids showed an aberrant morphology, whereas in HMW HA or Matrigel/collagen alone, significantly more spheroids appeared circular with a clear lumen (1% and 2% with aberrant morphology, respectively) (**Figures 8C,D**). To further study the effects of HMW HA on cholangiocyte growth, we cultured human cholangiocytes in a bile duct-on-a-chip microfluidic device [27] with an ECM of collagen alone or collagen with HMW HA. Initially, the cholangiocytes in both conditions formed a monolayer. This was followed, however, by the formation of outpouchings exclusively in the HMW HA ECM (**Figure 8E, leftmost panels**). This phenomenon was paralleled by an increase in cholangiocyte proliferation (**Figure 8E, middle and rightmost panels**). To assess the influence of LMW and HMW HA on collagen production in the EHBD, we employed a 2D co-culture system. We seeded a mixture of cholangiocytes and mesenchymal cells derived from neonatal mouse EHBDs and assessed collagen deposition and organization after treatment with media supplemented with water (control), LMW HA, or HMW HA (**Figure 8F**). After two weeks of culture, the monolayers comprised cholangiocyte clusters surrounded by dense layers of fibroblasts (**Figure 8F, leftmost panels**). SHG imaging showed that collagen deposition was increased by treatment with both LMW and HMW HA, but particularly by HMW HA (**Figure 8F, middle panels**). Collagen fibers were more aligned (higher peak around the 0° position) in the HMW HA-treated group compared to the control and LMW HA groups (p < 0.01 and p = 0.01, respectively) (**Figure 8F, rightmost panel**). Note that cholangiocytes were directly surrounded by HA suggesting HA production (**Figure S10**); in Figure S10, spheroids were cultured without HA. Together, these data show that HMW HA stimulates cholangiocyte spheroid formation and proliferation and suggest that it may provide a favorable environment for PBG organization in vivo. In contrast, LMW HA promotes aberrant spheroid morphology and impedes spheroid growth. This may have important implications for cholangiocyte structures surrounded by HA that becomes fragmented in pathological conditions. In addition, both LMW and HMW HA stimulate collagen production and alignment; this is essential for EHBD development as well as both regenerative (healing) and pathological (fibrotic) wound repair.

**Figure 8.**
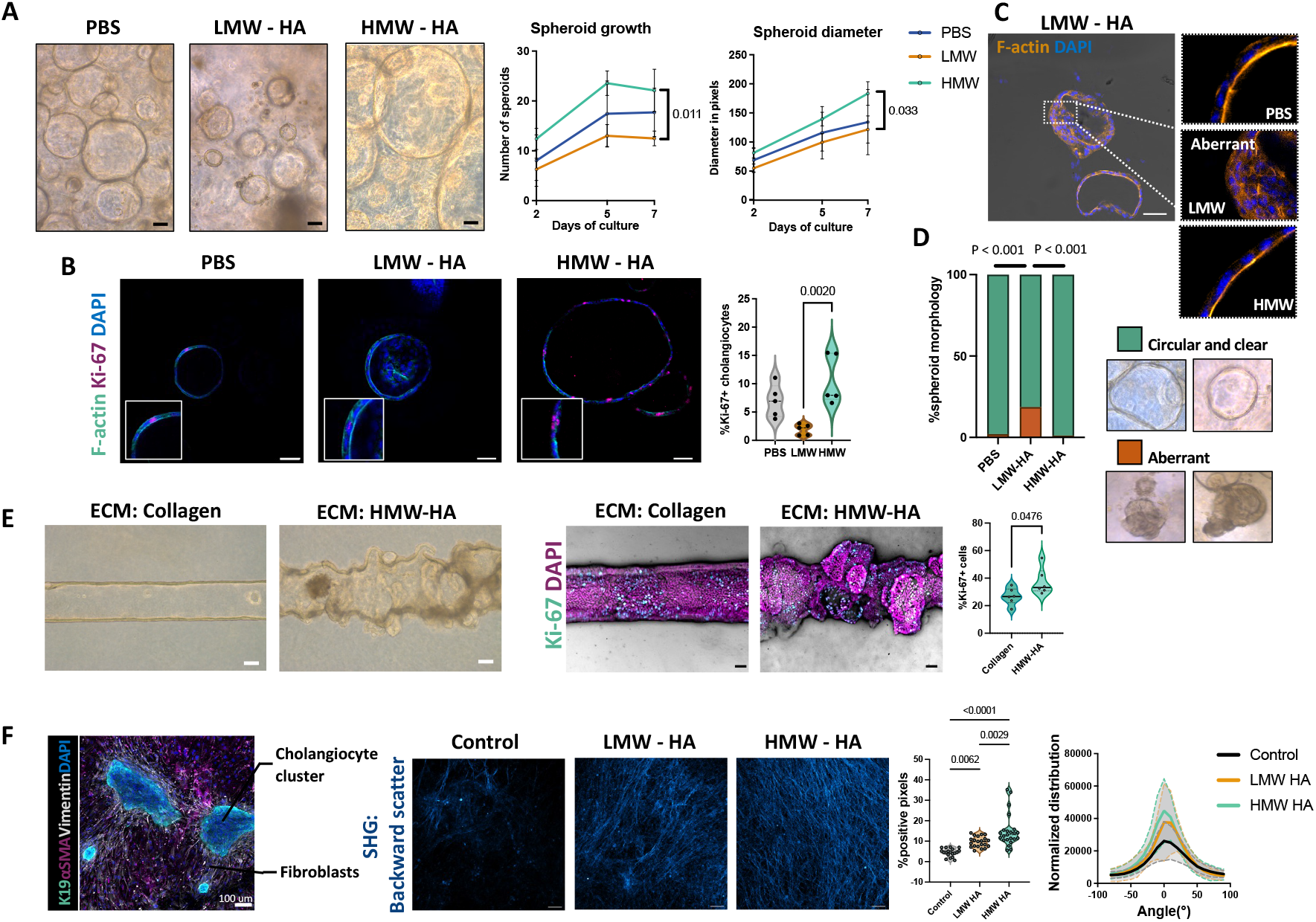
LMW and HMW have different effects on EHBD cholangiocytes and mesenchymal cells. **(A)** Spheroids derived from mouse primary cholangiocytes cultured in collagen with PBS, LMW HA, or HMW HA (n = 5 biological repeats with 2-3 technical replicates per condition). **(B)** Proliferation (Ki-67) of spheroids in each condition. **(C)** Spheroids stained with F-actin with bright field background. **(D)** Morphological scoring of spheroids in each condition. **(E)** Human cholangiocytes were seeded in the bile duct-on-a-chip device with a mixture of collagen alone or HMW HA with collagen in the ECM compartment. Immunofluorescence for Ki-67 on the right (n = 3 biological repeats with 2 technical replicates per condition). **(F)** (left) K19, aSMA, and vimentin staining of 2D cocultures. (middle) SHG imaging (back scatter) of the cholangiocyte/fibroblast-derived matrices in each condition (n = 3 biological repeats with 3 technical replicates per condition). Significance determined by area under the curve (A), Student’s t-test (B, E), Chi square test (D) and One-way and Two-way ANOVA for SHG intensity and collagen alignment, respectively (F). Graphs show mean ± SD (all panels). All scale bars = 50 μm.

## DISCUSSION

We demonstrate here that fetal EHBD injury leads to a vigorous program of fetal wound healing. Specifically, 1) the fetal/neonatal EHBD had high levels of HA compared to the adult EHBD; 2) in the fetus and neonate, HA was localized circumferentially immediately adjacent to the lumen, with a relatively thin layer of collagen at the periphery of the duct, while in the adult the collagen layer reached from the periphery to just below the mucosal layer; 3) EHBD injury in fetal sheep and in human BA resulted in an expansion of the HA layer encircling the bile duct lumen, with lumen narrowing and potential compression of the peripheral collagen layer; 4) sheep fetuses with EHBD injury demonstrated elevated bilirubin levels, consistent with EHBD obstruction, although no liver or EHBD fibrosis was noted; 5) the molecular weight of HA was significantly higher in the EHBDs of neonates compared to adults, and HA degradation in the neonate increased after injury; 6) LMW HA in 3D in vitro systems promoted an aberrant epithelial architecture, whereas HMW HA stimulates proliferation and the formation of PBG-like structures; and 7) both LMW and HMW HA stimulated collagen production in co-cultures of EHBD-derived fibroblasts and cholangiocytes. Collectively, these data show that fetal EHBD injury stimulates a fetal-type wound healing response but also suggest that the increased deposition of HA, while enhancing regeneration, leads to swelling and obstruction – an example of a beneficial response (fetal wound healing) “going bad”, and a new mechanism that may contribute to the pathophysiology of BA.

Fetal wound healing is characterized by regenerative, scarless repair. This pro-regenerative microenvironment features high levels of HMW HA and collagen-3 instead of the pro-fibrotic LMW HA and collagen-1 seen in scarring adult wounds [9,12,21]. Fetal wound healing across tissues – including lung, heart, tendon, and skin – is characterized in rodent and sheep models by prolonged increases in HA levels compared to adults, lasting up to three weeks after the initial injury [28]. In our fetal sheep model, EHBD injury lasting as long as 21 days was accompanied by marked expansion of the HA layer and increases in bilirubin levels. Although we were unable to monitor the consequences of injury longer than 21 days, the nature of the fetal wound healing program in other tissues suggests that expansion of the EHBD HA layer and subsequent narrowing of the lumen (as described below) could persist for weeks after the initial injury, potentially aggravating cholestasis and thereby EHBD wall damage.

In humans, transition to a so-called adult program of wound healing typically begins in the third trimester of pregnancy. Adult wounds contain predominantly LMW HA [26] and in a fetal rabbit model of skin wounds, experimental fragmentation of HA with hyaluronidase was associated with more ‘adult-like’ scarring, defined by excessive collagen production and neovascularization [29]. In the work we report here, damage to the neonatal rat EHBD led to increased HA degradation, reflecting a transition to adult wound healing. We propose that (persistence of) fetal EHBD injury or a subsequent regenerative response in late gestation could lead to increased HA degradation, consistent with features of adult wound healing (**Figure S11**).

Our results suggest that LMW HA in the EHBD wall during the transitional period and during adult wound healing could promote aberrant epithelial morphology and decreased cholangiocyte growth. The BA EHBD remnants (from Kasai surgeries) we examined had features of both fetal and adult wound healing, with an expanded peri-epithelial HA layer and epithelial structures that were phenotypically immature and morphologically distinct from normal PBGs and/or surface epithelium, consistent with fetal wound healing, but collagen organization reflecting an adult pathological process, as would be expected based on the age of BA patients at Kasai. The remarkable distribution of HA and collagen in both fetal EHBDs and BA remnants suggests that BA should be viewed in light of age-appropriate fetal-adult wound healing programs and that drawing parallels between BA and adult fibrosing cholangiopathies should be undertaken with caution.

The high concentrations of HA in the fetal EHBD may have implications beyond regenerative healing. An HA-rich ECM is a porous meshwork that exerts pressure on its surroundings due to repulsion between and within molecules [30]. Furthermore, owing to its hygroscopic properties, HA can swell up to 1,000 to 10,000 times, thereby shaping an environment in which cells and cell structures are distanced from each other, avoiding inhibitory contacts [21,31]. This malleable HA-rich ECM is ideal for cell migration, which may explain why PBGs are distributed evenly throughout the fetal EHBD but are in collagen-encircled clusters in the adult EHBD. This may also explain the formation of outpouchings in the collagen/HA-filled bile duct-on-a-chip – PBGs can easily increase in number in an HA-rich environment, as demonstrated by the remarkable PBG hyperplasia observed in fetal EHBDs.

The increased PBG volume and expanded HA layer during epithelial regeneration make it likely that fetal EHBDs can swell and are particularly susceptible to obstruction, potentially even during normal development. On the other hand, it is also possible that in the setting of EHBD injury during gestation, some ducts progress to total obstruction and BA but others undergo successful repair, potentially accompanied by transient lumen narrowing and bilirubin elevations. A recently-published large clinical trial which examined the efficacy of serum bilirubin measurements as diagnostic tools for BA reported that, of 123,279 total newborns tested, all of the 7 babies who went on to develop BA had elevated bilirubin levels at 60 h and two weeks after birth, but so did a significant number of newborns who never developed cholestatic symptoms or disease [1]. We speculate that these newborns experienced transient EHBD swelling causing elevated bilirubin levels either as part of a healing injury – a forme fruste of BA – or even as part of normal growth.

Steroid treatment of certain tumors leads to a reduction in HA synthesis and stromal edema [32,33]. Interestingly, the administration of steroids after a Kasai procedure did not influence post-operative survival or the number of transplantation-free months, but it did lead to decreases in serum bilirubin levels [34]. It is possible that steroids decreased HA synthesis and/or swelling in the large intrahepatic bile ducts of these patients, effectively widening the lumen. Notably, this effect was most prominent in the younger cohort, the group that would be most likely to have high levels of HA. It is conceivable that as BA progresses, more collagen is deposited around the ducts and other epithelial structures, limiting the effect of steroids on reducing EHBD wall volume.

We observed numerous goblet cells in the EHBDs of the hypoxic animals, although intestinal metaplasia was found in some control fetal sheep EHBDs as well. The occurrence of goblet cells in the surface epithelium of the EHBD has been described previously in autopsy samples from human newborns between 28-41 weeks of gestation [22], although it is not a general feature of BA. We noted higher overall viscosity of fetal as compared to adult bile in sheep and it is possible that the same is true in humans; we hypothesize that the combination of swelling and luminal narrowing from HA and viscous bile contributes to luminal obstruction and BA pathophysiology, although this would be difficult to test in human fetuses and neonates.

In summary, the damage-repair response of the fetal EHBD is an example of fetal wound healing. The expansion of the HA layer in the EHBD wall potentially leads to obstruction of the EHBD lumen, which can be detected by elevated serum bilirubin levels. The initial lumen narrowing could lead to progression of fibrosis/scarring when accompanied by the transition to an adult-type wound healing in late gestation, including increased degradation of HA. Viewing the damage-repair response of fetal EHBDs in context of fetal wound healing will be crucial to understanding the early pathogenesis of BA, including rapid progression to biliary fibrosis after birth, and to the development of potential treatments.

## ABBREVIATIONS

AE2: anion exchange protein 2
Alc-B: alcian blue
BA: biliary atresia
DOL: day of life
EHBD: extrahepatic bile duct
ECM: extracellular matrix
FFPE: Formalin-fixed paraffin embedded
GA: gestational age
HA: hyaluronic acid
HAS: hyaluronic acid synthase
HMW: high molecular weight
K19: keratin 19
LMW: low molecular weight
PAS: periodic acid Schiff
PBG: peribiliary gland
PBS: phosphate buffered saline
PDX1: pancreatic and duodenal homeobox 1
PSC: primary sclerosing cholangitis
SHG: second harmonic generation

## ACKNOWLEDGEMENTS

We are thankful to the UPenn Cell and Developmental Biology Microscopy Core, the Penn Vet Imaging Core, and the NIDDK Center for Molecular Studies in Digestive and Liver Diseases Molecular Pathology and Imaging Core and Host Microbial Analytic and Repository Core (NIH P30DK050305).

