## Supplemental methods and figures for "Injury and a program of fetal wound healing in the fetal and neonatal extrahepatic bile duct"

Figure S9…………………………………………………………………………………19-20

Figure S10………………………………………………………………………………..21

Figure S11………………………………………………………………………………...22

Supplementary references…………………………………………………………...……23

### SUPPLEMENTARY METHODS AND MATERIALS

#### Histochemistry and immunostaining

All bile duct sections were fixed in 10% formalin, embedded in paraffin, and cut into 5 μm sections. Sections for Sirius Red, periodic acid Schiff (PAS), Alcian Blue (Alc-B), and Hematoxylin and Eosin (H&E) were processed according to standard protocols. Antibodies used for immunostaining are listed in **Table S1**. Paraffin-embedded sections were deparaffinized with xylene and rehydrated through a graded series of alcohol and distilled water. Antigen retrieval was performed in 10 mM citric acid buffer (pH 6.0). For diaminobenzidine (DAB) labelling, sections were incubated with 3% $H_2O_2$ to quench endogenous peroxidases and blocked with StartingBlock™ T20/phosphate buffered saline (PBS) Blocking Buffer (Thermo Fisher Scientific, Waltham, MA, USA) and Avidin D and Biotin Blocking Reagents, prior to incubation with primary antibodies overnight at 4°C. The next day, sections were incubated with secondary antibodies at a concentration of 1:200 for 30 minutes at 37°C and visualized using an Avidin-Biotin Complex detection system (Vector Elite Kit, Vector Laboratories, Burlingame, CA, USA). Signals were developed by a DAB substrate kit for peroxidases (Vector Laboratories) and counterstained with hematoxylin. For immunofluorescence, sections were blocked with 5% bovine serum albumin and permeabilized with 0.4% Triton X-100 prior to antibody incubation. Cy2-, Cy3-, and Cy5-conjugated secondary antibodies were used to visualize the marker (1:500, Vector Laboratories). For the HA binding protein assay, binding protein to HA was already biotinylated and, therefore, the secondary antibody step for DAB labelling was omitted from the protocol.

#### Second harmonic generation (SHG) imaging

Fixed sections were imaged using a Leica SP8 confocal/multiphoton microscope and Coherent Chameleon Vision II Ti:Sapphire laser (Leica, Buffalo Grove, IL, USA) tuned to a wavelength of 910 nm. SHG generated both backward and forward scatter in the blue and red channel, respectively. The green channel was used for concomitant staining with HA.

**Image analysis**

Image analysis of stained sections was performed with Fiji ImageJ and QuPath v0.2.0 software [1]. QuPath was used to manually select of areas of interest within images (e.g., PBGs, surface epithelium, stroma, entire EHBD wall, spheroids). PBG area was calculated by measuring % area occupied by PBGs of the total EHBD wall area. The Pixel classification or Positive cell detection tool was used to identify the positive area or positive cells respectively within the Region of Interest. The directionality of collagen in 2D cell cultures was calculated from SHG signals with the ImageJ plugin OrientationJ. The intensity of immunostaining in the 2D cell cultures was quantified with the threshold and % area functions of ImageJ. Measurements of collagen end points, length, density, and curvature were carried out using the ImageJ plugin TWOMBLI [2] after 250 μm x 250 μm areas without vascular, neural, or epithelial structures were selected using the grid function of QuPath. Curvature was defined as mean change in angle moving incrementally along fibers of 90 μm (the window was set manually by measuring fibers in the image). Endpoints were defined as the number of collagen fiber endpoints in a certain image. The normalized value was obtained by dividing the raw values by the total length of fibers in the mask. Fiber length was calculated from the fiber endpoints and branch points [2]. Fiber density was calculated as %positive SHG pixels per image. All analyses were performed in a blinded fashion.

**Bilirubin oxidase treatment**

After deparaffination and rehydration of FFPE sections, tissue was treated with 25 U/mL bilirubin oxidase at 37°C for 6 days and rinsed with PBS prior to PAS staining.

**Primary cholangiocyte isolation and 2D culture**

Primary neonatal extrahepatic cholangiocytes and mesenchymal cells were isolated by outgrowth from 2–3-day old BALB/c mouse pups in mouse cholangiocyte media as described previously [3]. 4-5 neonatal EHBDs were required to obtain ~5x$10^6$ cells at ~70% confluency. For 2D cultures, cholangiocytes and mesenchymal cells were counted and plated on glass coverslips that were incubated for at least 4 h with mouse cholangiocyte media (components are described in [3]) prior to seeding. Cells were cultured in mouse cholangiocyte media supplemented with 0.1 mg/ml ascorbic acid and, from day 2 onwards, with di-$H_2O$, 0.5 mg/ml LMW HA (60kDa), or 0.5 mg/ml HMW HA (1.5MDa) (LifeCore Biomedical, Chaska, MN, USA). Media was changed and re-supplemented every two days. Cells in monolayers were fixed on day 14 for further processing.

**Tissue homogenization and HA and collagen quantification**

Rat and mouse EHBDs were weighed and homogenized in PBS using a Bead Mill homogenizer. Homogenates were lysed by adding RIPA buffer and dissolved protease inhibitor cocktail tablets (cOmplete Protease Inhibitor Cocktail; Roche, Indianapolis, IN, USA). Aliquots were stored at -20°C until further processing. HA amounts were measured with the Hyaluronan Duoset ELISA Kit (R&D systems, Minneapolis, MN, USA) according to the manufacturer's instructions. To prepare rat and mouse EHBD samples for collagen quantification, they were hydrolyzed with 12M HCl at 120°C for 3 h after which 50 μL was evaporated in a 60°C oven. Collagen was quantified with the Hydroxyproline Assay Kit from Sigma Aldrich (Burlington, MA, USA) according to the manufacturer's instructions.

**HA isolation for size measurements**

HA was isolated from EHBD homogenates (dissolved in RIPA buffer and inhibitor cocktail tablets as described above) following general protocols reported previously [4] with minor modification. First, proteins were digested using proteinase K (Ref. AM2548, Invitrogen) following the manufacturer's directions. Phenol:chloroform:isoamyl alcohol, 25:24:1 (Ref.

327111000, Thermo Fisher Scientific) was added to the sample at an equal volume and partitioned into the aqueous phase (containing HA, nucleic acids, and other polysaccharides) and organic phase (containing proteins and other debris) in a phase-lock gel tube (Ref. 2302830, QuantaBio) by thorough mixing and centrifugation (14,000 g for 15 min at 20°C). The same process was repeated twice with pure chloroform (Ref. AC423555000, Thermo Fisher Scientific) to remove residual phenol. Next, biomagnetic precipitation was employed to capture HA from the solvent-extracted aqueous samples. For this, superparamagnetic beads (Dynabeads™ M-280 Streptavidin, Ref. 11206D, Thermo Fisher Scientific) were washed according to the manufacturer's directions and then incubated (room temperature for 1 h under agitation) in 1X PBS with biotinylated versican G1 domain (bVG1, Ref. G-HA02, Echelon Biosciences) at a ratio of 1 µg bVG1 per 100 µg beads. After thorough washing and resuspension of the bVG1-conjugated beads to a concentration of 10 mg/mL in 1X PBS, beads were separated in 150 µL aliquots. Aqueous sample was added dropwise to avoid capture size bias and the mixture was incubated at room temperature for at least 2 h before removing unbound material and washing the beads. Finally, beads were incubated in 6M LiCl for 1 h to elute HA from the beads into solid-state nanopore (SSNP) measurement buffer.

**Solid-state nanopore analysis**

SSNPs consisting of a single pore (diameter 6-11 nm) in a silicon nitride membrane (20-30 nm thickness) were either fabricated using a Helium ion milling method reported previously[5] or produced commercially through conventional silicon processing methods (Norcada, Edmonton, Canada). For HA measurement [6], a chip was rinsed with ethanol and water, dried under filtered air flow, treated with air plasma (30 W, Harrick Plasma, Ithaca, NY) for 2 min per side, and mounted into a custom 3D printed flow cell (Carbon, Redwood City, CA). Measurement buffer (6M LiCl, 10 mM Tris, 1 mM EDTA, pH 8.0) was then introduced to contact each side of the membrane and Ag/AgCl electrodes were used connect to an Axopatch 200B patch-clamp amplifier (Molecular Devices, San Jose, CA).

Isolated HA suspended in measurement buffer was then introduced to one side of the SSNP and measurements were performed by applying either a 200 or 300 mV bias and monitoring trans-membrane current at a rate of 200 kHz through a 100 kHz four-pole Bessel filter. Data were collected using a custom LabVIEW program (National Instruments, Austin, TX) and analysis was performed with an additional 5 kHz low-pass filter applied. Molecular translocations were marked by temporary reductions in the ionic current (events), analyzed using thresholds described previously [7]. The magnitude of each event was mapped to a molecular weight using a calibration curve [6]. All data sets consisted of at least 500 individual events.

**3D cholangiocyte spheroid culture**

After reaching ~70% confluency, primary neonatal mouse extrahepatic cholangiocytes and mesenchymal cell cultures were isolated as previously described [3] and divided into three equal suspensions, one for each matrix mixture. Cell pellets were resuspended in a mixture of collagen/Matrigel with PBS, LMW HA or HMW HA, with final concentrations of: 1.25 mg/ml rat tail collagen I (Ibidi, Fitchburg, WI, USA), 1 mg/ml HA, and 30% Matrigel (Corning, Corning, NY, USA). Droplets of 200 μL cell/gel mixtures were pipetted into culture dishes with a glass bottom (MatTek, Corporation, Ashland, MA, USA), 2 for each condition. Mesenchymal cells were co-cultured with cholangiocyte spheroids to maintain mesenchymal-epithelial communication. On day 8, spheroids were fixed and stained for F-actin (1:400, phalloidin-tetramethylrhodamine B isothiocyanate; Thermofisher) or with Ki-67 (1:100) or were processed for histology (H&E).

**Bile duct-on-a-chip experiments**

Microfluidic devices were fabricated with polydimethylsiloxane (PDMS; Sylgard 184; Dow-Corning, Midland, MI, USA) as described previously [8]. A mixture of 2.5 mg/mL rat tail type 1 collagen (pH = 7.0, Thermofisher) with or without HMW HA (final concentration: 1 mg/ml) was used as the ECM compartment and polymerized for 20 minutes at 37°C with a needle of 200 μm diameter in situ to form the channel. After the needle was removed, the channel was washed three times with PBS and the ends were sealed with vacuum grease. Primary human cholangiocytes were isolated from fresh human bile duct tissue, obtained post-mortem from healthy individuals (HPPA program, see 'human samples' for protocol number), followed by cell expansion in an organoid culture system as previously described [9]. Cholangiocytes were dissociated to single cell suspensions before they were seeded in the channels. A suspension of $0.5x10^6$ cholangiocytes/mL was introduced through the reservoir ports (40 μl to the left port and 30 μl to the right port) and allowed to adhere to the top surface of the channel in an inverted position and to the bottom surface for 5 minutes to ensure an even distribution of the cholangiocytes in the channel, followed by removal of unattached cells by rinsing with cell culture media (William's E+ medium supplemented with 500 ng/ml R-spondin, 50 ng/ml EGF, 100 ng/ml DKK-1 and 10 μM Y27632). After two days of culture in this media, cells were cultured in William's E+ medium with 500 ng/ml R-spondin, 50 ng/ml EGF and 100 ng/ml DKK-1 until a compact monolayer was formed. Afterwards, cells were cultured for approximately another week prior to fixing and processing for immunohistochemistry with Ki-67 (1:100).

**Whole-duct explant culture**

Intact EHBDs were isolated from 4-to-5-day-old Sprague-Dawley rats and maintained in ice-cold cholangiocyte media [3] until being placed in glass vials that allowed for air exchange. The glass vials were filled with 2 mL mouse cholangiocyte media without fetal bovine serum and supplemented with 2 μg/mL biliatresone in DMSO [10] or an equivalent amount of DMSO alone (0.0002%). The vials were placed in a Vitron Dynamic Organ Culture Incubator and maintained at 37°C in 95% $O_2$/5% $CO_2$ with rotation for 5 days. The media was changed daily. After 5 days of incubation, the ducts were snap frozen in O.C.T compound (Tissue Tek, VWR, Bridgeport, NJ, USA) and sectioned into 5 μm thick sections.

**Hyaluronic acid in situ zymography**

Snap frozen EHBD sections were adjusted to room temperature for 10 min and rinsed in PBS. Next, sections were incubated with 0.1 mg/ml fluorescein-tagged HA (Sigma Aldrich) overnight at 37°C. Hyaluronidase (from bovine testes, Sigma Aldrich) was added to 0.5 mg/mL as positive control. Other control sections were incubated for 15 minutes and 2 hours. Sections were counterstained with DAPI and mounted before imaging.

**Statistical analysis**

Continuous data are presented as mean ± standard deviation (SD). The normality of data was first evaluated by using the Shapiro-Wilk test and visualizing the QQ-plots. Normally-distributed data were analyzed using the Student's t test and non-parametric data with the Mann-Whitney test. We used one-way ANOVA followed by Tukey's post hoc analysis to compare more than 2 groups and two-way ANOVA when comparing multiple independent variables. Frequencies of stratified vs. non-stratified spheroids were compared using the Chi Square test. Differences in spheroid growth and diameter over time were analyzed by calculating the p-value from the area under the curve (AUC) and the standard error (SE). The Spearman non-parametric correlation test was used to determine the relationship between two variables. Tests used for each comparison are detailed in the figure legends; p-values are specified in the graphs when statistically significant. A p-value $<0.05$ was considered statistically significant. Analyses were performed using Graphpad Prism v9.0 (Graphpad Software, San Diego, CA, USA).

### SUPPLEMENTARY TABLES

#### Table S1. List of antibodies used for (immuno-)stainings.

| Antigen | Used as marker for | Host | Supplier | Catalog. No. | Dilution |
|---|---|---|---|---|---|
| AE2 | Mature cholangiocytes – bicarbonate transporter | Mouse | Santa Cruz | Sc-376632 | 1:50 |
| $\alpha$-SMA | Activated fibroblasts | Mouse | Sigma Aldrich | A2547 | 1:100 |
| CK19 | Cholangiocytes | Rat | DSHB | Troma III, AB 2133570 | 1:50 |
| Collagen I | Collagen I | Goat | Southern Biotech | 1310-01 | 1:100 |
| HA | Hyaluronic acid | Biotinylated binding protein | EMD Millipore | 385911 | 1:100 |
| HAS1 | Hyaluronic acid production | Rabbit | Novus Biologicals | NBP2-39087 | 1:100 |
| HAS2 | Hyaluronic acid production | Mouse | Abcam | AB140671 | 1:100 |
| HYAL1 | Hyaluronic acid degradation enzyme | Rabbit | Thermofisher | PA5-51686 | 1:100 |
| HYAL2 | Hyaluronic acid degradation enzyme | Rabbit | Novus Biologicals | NBP1-81283 | 1:50 |
| Ki67 | Proliferation | Rabbit | Abcam | Ab14917 | 1:100 |
| PDX1 | Endoderm progenitor marker | Rabbit | Novus Biologicals | NBP2-22150 | 1:100 |
| Sox9 | Biliary progenitor cells | Rabbit | Millipore | AB5535 | 1:500 |
| Vimentin | Mesenchymal cells | Chicken | Novus Biologicals | NB300-223 | 1:1000 |

Abbreviations: AE2, anion exchanger 2; CK19, cytokeratin 19; HA, hyaluronic acid; HAS1/2, hyaluronic acid synthesis; HYAL1/2, hyaluronidase; PDX1, pancreatic and duodenal homeobox 1.

**Table S2. Total number of translocation events collected with each sample during solid-state nanopore analysis.**

| | | Number of ducts in homogenate | Translocation events |
|---|---|---|---|
| **Mouse** | **Neonate** | 15 | 1741 |
| | | 15 | 1316 |
| | | 16 | 1272 |
| | | 8 | 1218 |
| | | 10 | 1809 |
| | **Adult** | 3 | 2908 |
| | | 6 | 500 |
| | | 3 | 1231 |
| | | 4 | 1580 |
| **Rat** | **Neonate** | 5 | 5139 |
| | | 13 | 4183 |
| | | 11 | 1507 |
| | **Adult** | 2 | 1980 |
| | | 2 | 1995 |
| | | 5 | 1940 |

**SUPPLEMENTARY FIGURES**

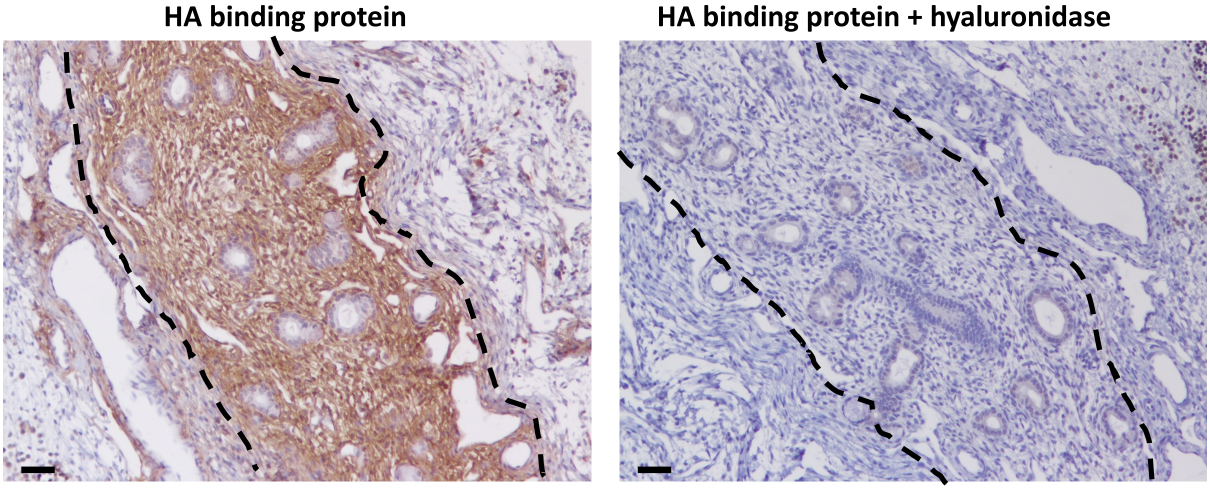

**Supplementary Figure 1. Confirmation of HA binding protein specificity.** Section on left untreated, on right treated with hyaluronidase at 37°C for 1 h; then incubated with HA binding protein overnight at 4°C. Scale bars = 50 μm.

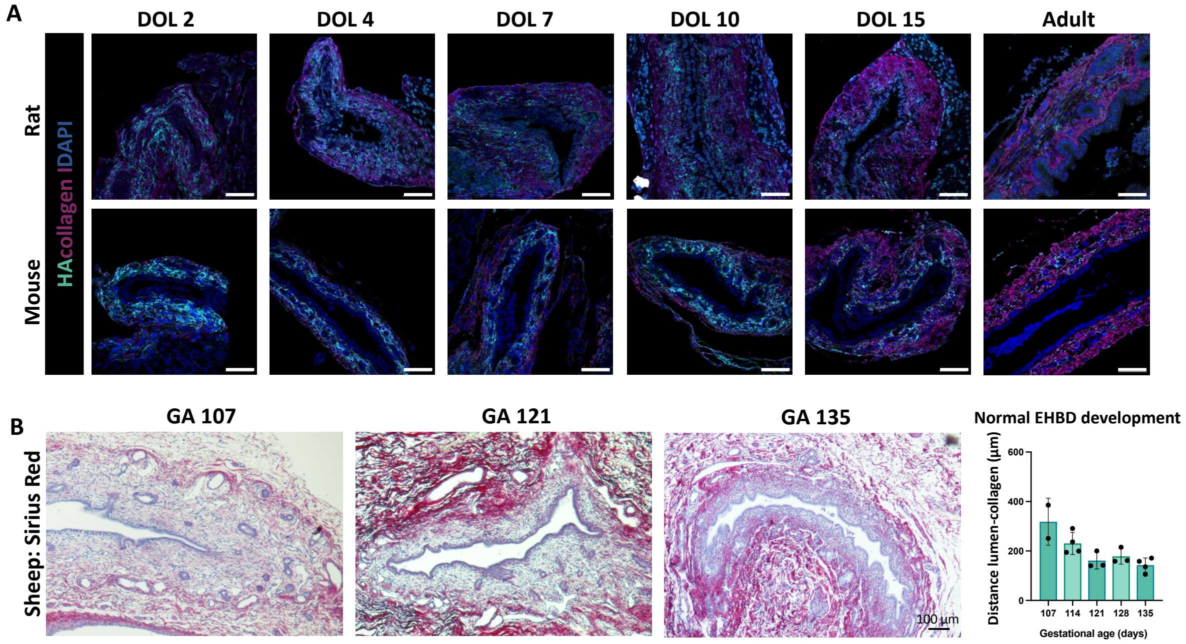

**Supplementary Figure 2. The major ECM components collagen and HA change during EHBD development. (A)** Representative images of mouse and rat EHBDs at postnatal day of life (DOL) 2, 4, 7, 10, 15 and at adulthood; n = 2-4 individuals for each species per time point. Staining for HA (cyan) and collagen I (magenta). DOL, day of life. Scale bars = 50 μm. **(B)** Representative images of Sirius Red staining of fetal sheep EHBDs at gestational ages 107, 121, and 135 days. Data are shown as mean ± SD distance in μm from the lumen to the collagen layer for n ≥ 2 individuals per time point. ≥ 5 measurements were made per image. No statistics were performed on the data due to the low number of samples in the baseline group (n=2). EHBD, extrahepatic bile duct; GA, gestational age.

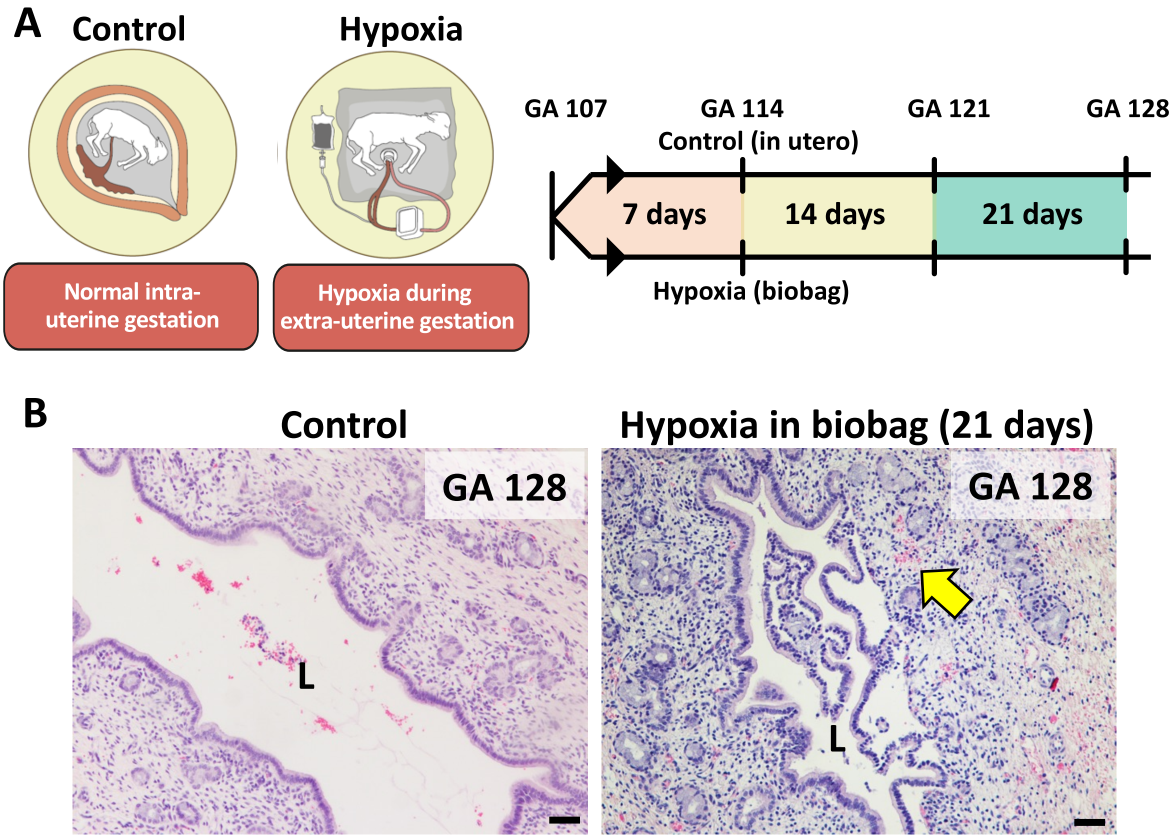

**Supplementary Figure 3. Model of prenatal hypoxia of fetal sheep in an external biobag. (A)** Experimental setup of the fetal sheep hypoxia model. We used an extra-uterine life support system to manipulate gas exchange in a controlled setting. The animals in the control group remained in utero (where average oxygen levels are 20-25 ml/kg/min) whereas the hypoxia group in the biobag was supplied with lower oxygen levels (average oxygen levels 14-16 ml/kg/min). Animals were placed into the external device at a gestational age of ~107 days and euthanized after 7, 14, and 21 days (gestational ages of 114, 121, and 128 days). GA, gestational age. **(B)** Typical EHBD histology of fetal sheep at a gestational age of 128 days (n=4) and of fetal sheep at corresponding age, but maintained in hypoxic conditions for 21 days (n=4). Arrow points to mural bleeding. GA, gestational age; L, lumen. Scale bars = 50 µm.

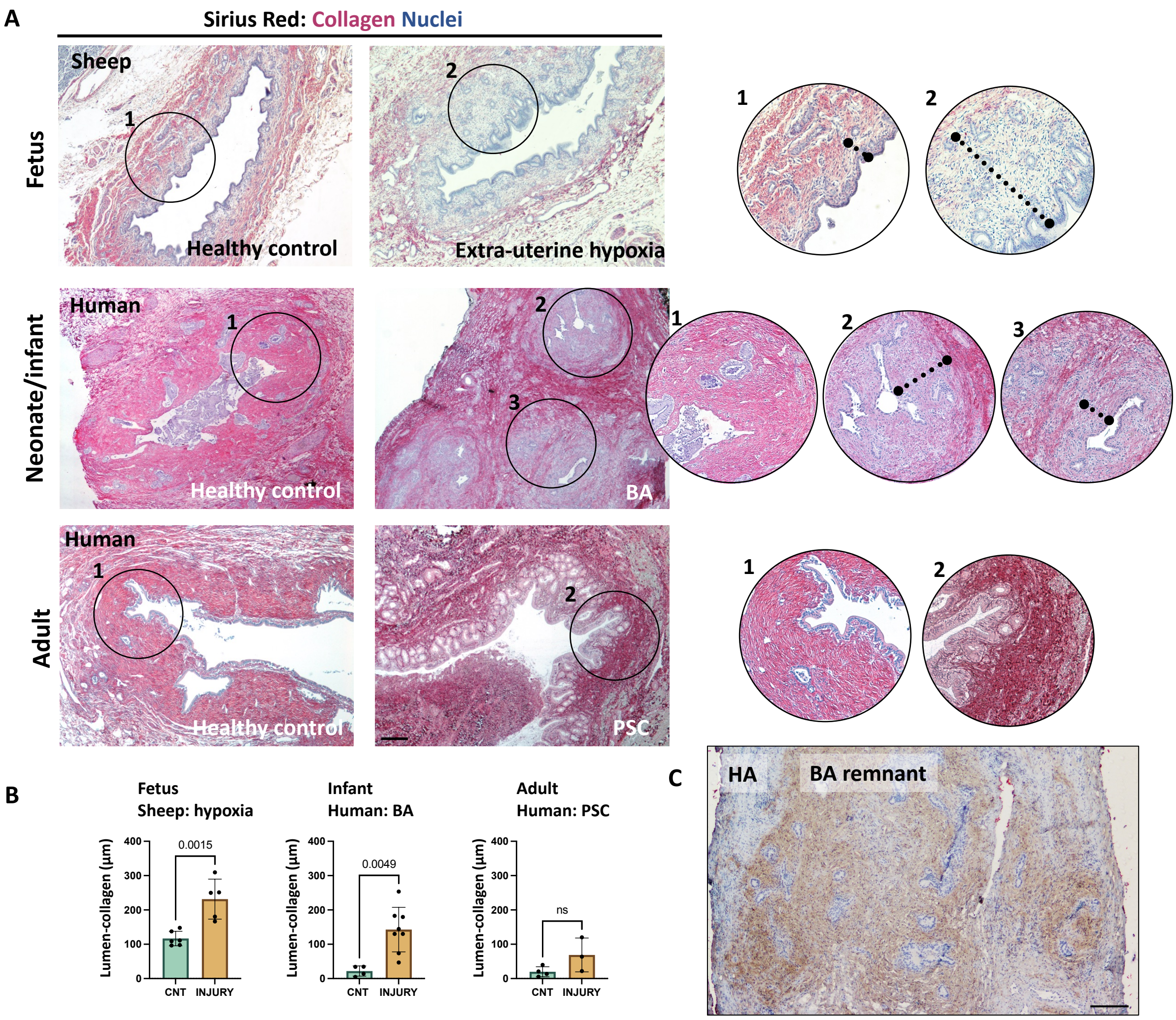

**Supplementary Figure 4. Distance between the lumen and the collagen layer is greater in damaged compared to control fetal and neonatal EHBD. (A)** Representative images of Sirius Red staining of damaged EHBD sections from sheep fetuses, human neonate and infant, and human adults and EHBD sections from age-matched subjects. Circles are shown at higher magnification on the right, as numbered. Dashed lines show distance measurements. PSC, primary sclerosing cholangitis. Scale bar = 200 µm. **(B)** Data presented as mean ± SD distance in µm from the lumen to the collagen layer. n = 3-8 individuals. For each image, ≥ 5 measurements made. Significance determined by Student's t test. BA, biliary atresia; CNT, control; ns, non-significant; PSC, primary sclerosing cholangitis. **(C)** HA staining of BA remnant typical of 14 tested. Scale bar = 200 µm.

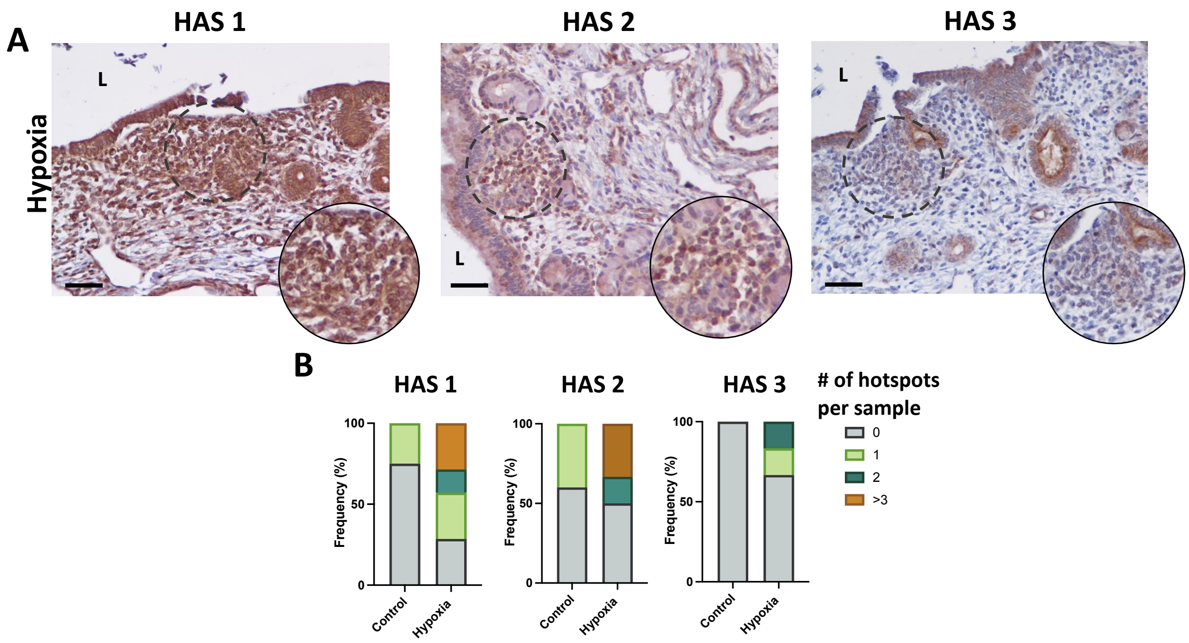

**Supplementary Figure 5. Prenatal hypoxia of the EHBD causes accumulation of HAS1-3 expressing mesenchymal cells. (A)** Immunohistochemistry for HAS1-3 in representative EHBD sections from fetal sheep in the hypoxia group. ‘Hotspots’, defined as clusters of HAS1-3+ mesenchymal cells, are shown in the magnified boxes. HAS, hyaluronic acid synthases. **(B)** Graphs represent the proportion of sections in which the indicated number of hotspots in control and hypoxic fetal sheep EHBDs were observed. Hotspots were identified based on recognition of the investigator, and analyses were done in a blinded fashion. n = 4-7 sections in each group.

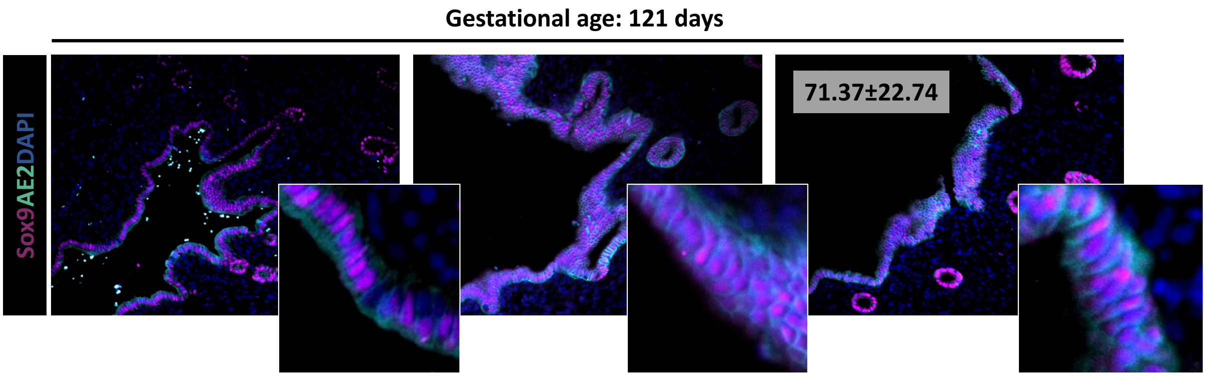

**Supplementary Figure 6. Sox9 is expressed in the fetal surface epithelium.**

Representative images of immunostains for the endoderm progenitor cell marker Sox9 (magenta) and the bicarbonate transporter AE2 (cyan) of normal fetal sheep at a gestational age of 121 days. Data shown in the right image represent mean ± SD of %Sox9+ cholangiocytes in the surface epithelium of n = 3 normal fetal sheep EHBDs at gestational age 121 days. Original images x20.

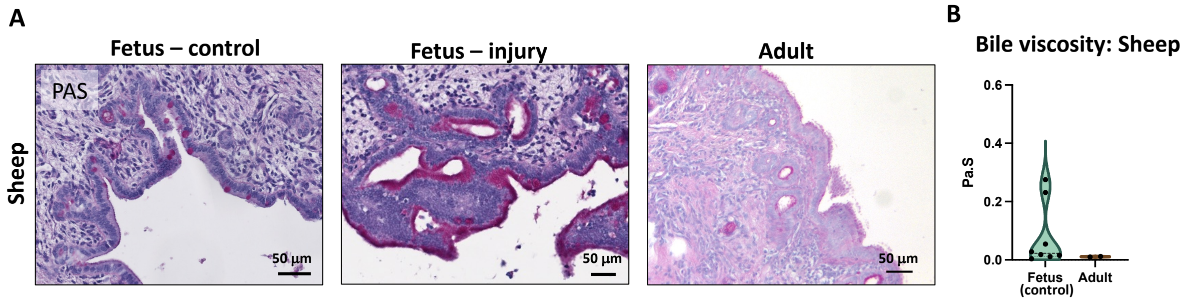

**Supplementary Figure 7. Increased mucin-containing cells and bile viscosity in fetal EHBDs (A)** Periodic acid Schiff (PAS) staining of samples from the EHBDs of a normal and an injured sheep fetus at gestational age 121 days, and the EHBD of an adult sheep. $n \geq 3$ for each group. **(B)** Bile viscosity at 25°C in Pa.sec from 40 μL bile samples of normal sheep fetuses at gestational ages of 114, 121, 128, and 135 days vs. bile of two adult sheep. Values of shear rates between 0.5 and 5 / s were selected to filter out noise. No statistics were performed on the data due to a low number of samples in the adult group (n=2).

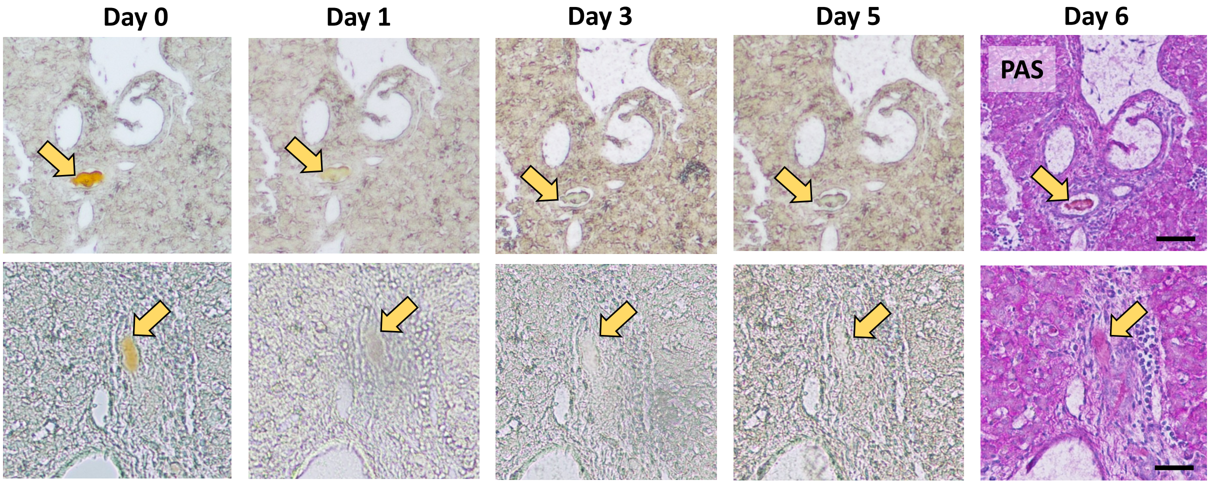

**Supplementary Figure 8. Bilirubin oxidase treatment removed the yellow color of small intrahepatic bile plugs.** Representative images of fixed liver sections from fetal sheep in the hypoxia group containing bile plugs (n = 3). Sections were treated for 6 days with 25 units/ mL bilirubin oxidase at 37°C. Images were taken daily during bilirubin oxidase treatment to demonstrate removal of pigment. Arrows point to bile plugs stained after 6 days of treatment with periodic acid Schiff (PAS). Scale bars = 50 μm.

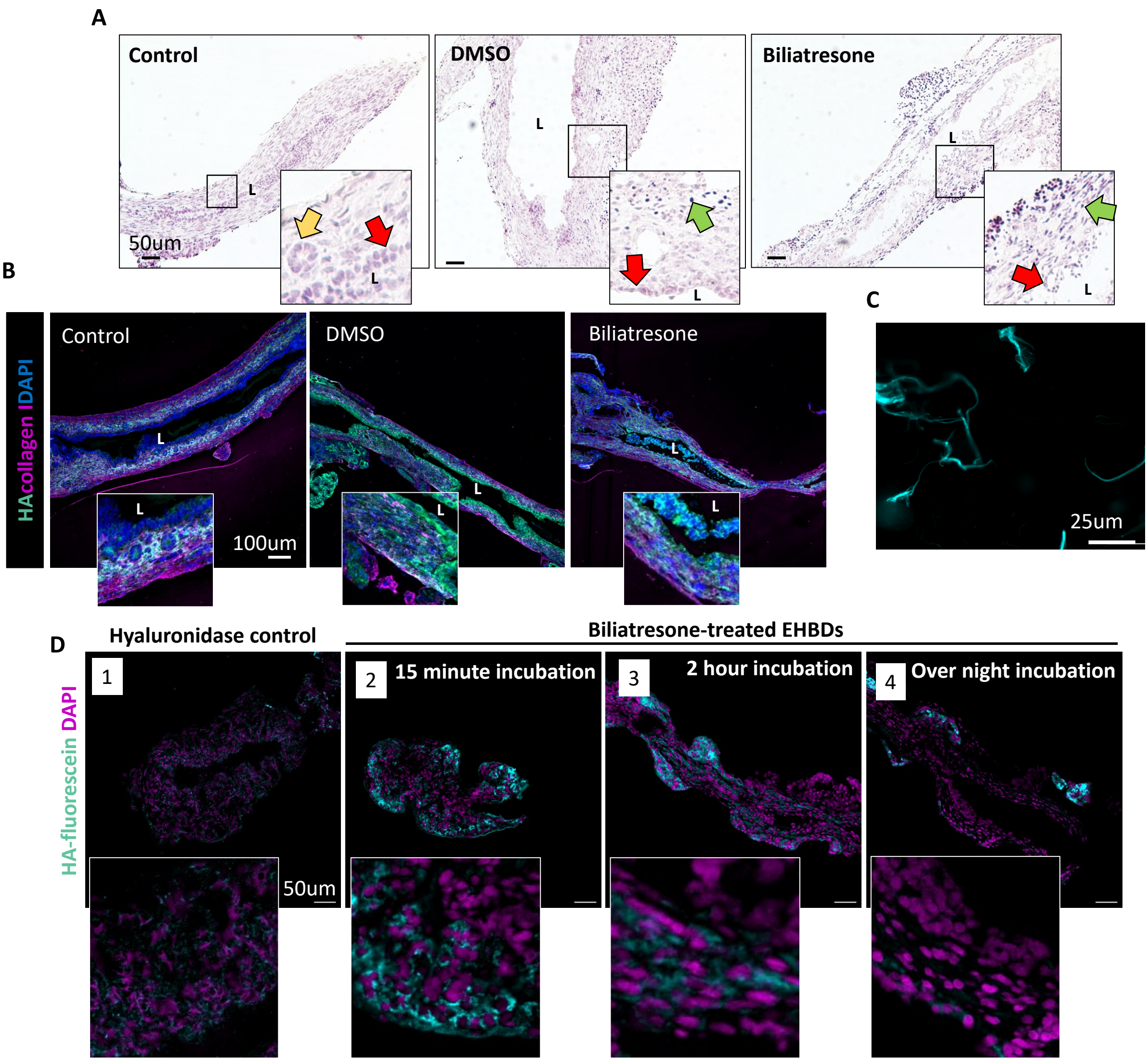

**Supplementary Figure 9. Treatment with biliatresone caused damage to rat neonatal EHBDs in a whole explant culture system. (A)** H&E-stained images from snap-frozen rat EHBDs after harvest at DOL 4/5 (control) or harvested and treated with DMSO or biliatresone for an additional 5 days in the high oxygen incubator, corresponding to those in Figure 6C. (**Left**) The control samples showed intact PBGs (yellow arrow), single or double layered surface epithelium (red arrow) and open lumens. The mesenchyme appeared viable with sporadically some pyknotic cells (indicating cell death) located on the far-outside of the ducts. (**Middle**) EHBDs that were treated with DMSO presented typically with intact single or double layered surface epithelium (red arrow), albeit occasionally appearing flat (indicating newly regenerated epithelium). Pyknotic cells were located throughout the duct wall, but predominantly in the outer layers (green arrow). (**Right**) Epithelium in the biliatresone-treated EHBDs was sloughed off (red arrow) and mesenchyme showed increased pyknotic cells compared to the other groups (green arrow); these were predominantly located in the outer layers but cell death in the subepithelial layer was not uncommon. Scale bars = 50 μm. **(B)** Representative images of stainings for HA and collagen I of neonatal rat EHBDs that have been fixed immediately after harvest at DOL 4/5 (control) or harvested and treated with DMSO or biliatresone for an additional 5 days in the high oxygen incubator. (**Left**) The control samples showed the typical HA-collagen organization as could be expected based on age (4/5 day old rat neonates). (**Middle and Right**) The DMSO and biliatresone treated EHBDs showed a mixed collagen-HA ECM with HABP staining the cytoplasm of cholangiocytes. Scale bars = 100 μm. **(C)** Visualization of structure of fluorescein-tagged HA before application on slides. **(D)** Controls for in situ zymography. A control sample was incubated with hyaluronidase together with the fluorescein tagged HA overnight (1) showing quenching of the signal; biliatresone-treated samples were incubated for 15 minutes total (2), or 2 h (3) showing still some signal, indicating incomplete degradation. A biliatresone-treated sample incubated overnight showing complete quenching of the signal (4). Scale bars = 50 μm.

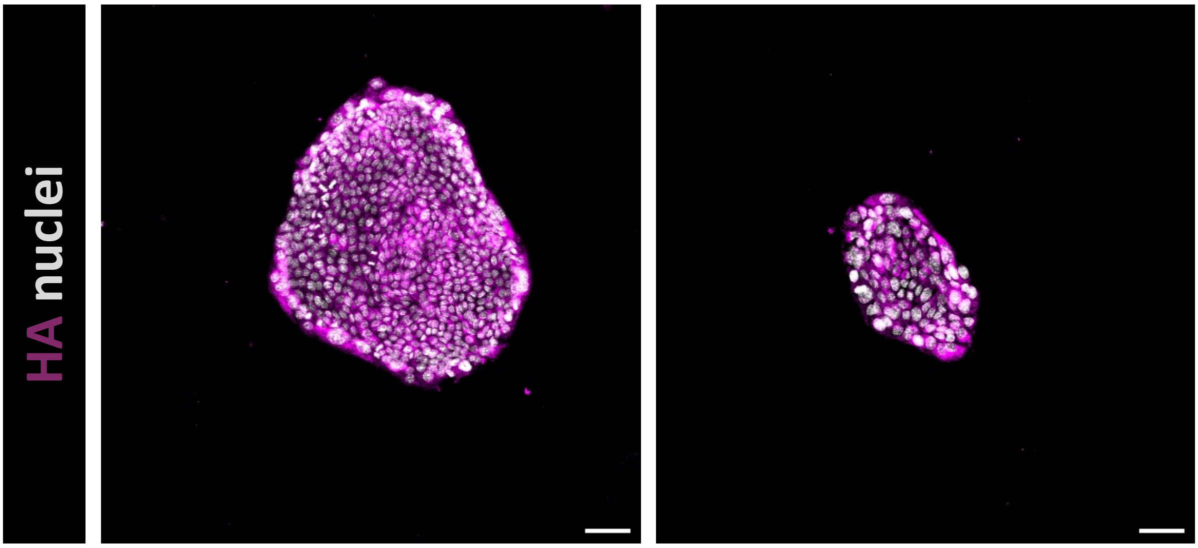

**Supplementary Figure 10. Neonatal mouse cholangiocyte spheroids produce HA during growth.** Representative images of spheroids after staining with HA binding protein (n = 3). Spheroids were cultured in a collagen/Matrigel mixture, without HA. Scale bars = 50 μm.

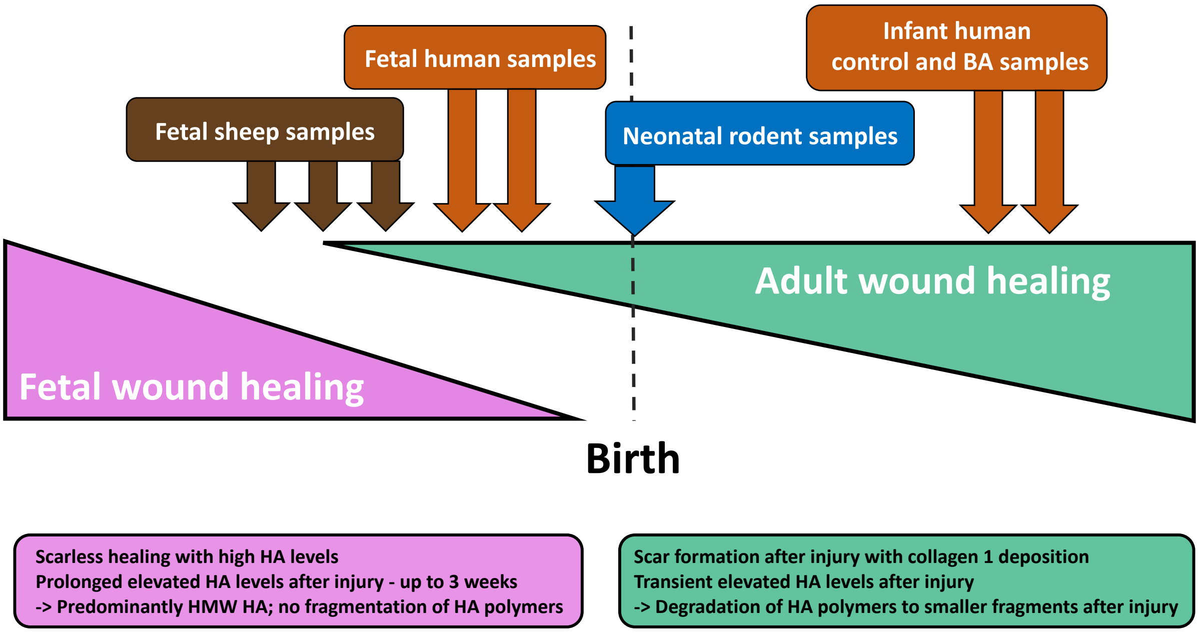

**Supplementary Figure 11. Schematic representation of features of fetal and adult wound healing before and after birth.** EHBD samples that were used in this study are indicated on the timeline. Gestation ages of the fetal sheep samples were 107, 114, 121, 128, and 135 days of 147 total (i.e., full term); corresponding to late second and third trimester. Gestation ages of fetal human samples were 22, 34, 36, and 39 weeks of 39 weeks total (i.e., full term) corresponding to the third trimester. HMW HA, high molecular weight HA.
